# Architecture and regulation of filamentous human cystathionine beta-synthase

**DOI:** 10.1101/2023.02.15.528523

**Authors:** Thomas J. McCorvie, Henry J. Bailey, Claire Strain-Damerell, Arnaud Baslé, Wyatt W. Yue

**Affiliations:** Centre for Medicines Discovery, Nuffield Department of Clinical Medicine, University of Oxford, Oxford OX3 7DQ, UK; Biosciences Institute, The Medical School, Newcastle University, Newcastle upon Tyne, NE2 4HH, UK; Institute of Biochemistry II, Faculty of Medicine, Goethe University Frankfurt, Frankfurt, Germany; Research Complex at Harwell, Harwell Science and Innovation Campus, Didcot, OX11 0FA, UK

**Keywords:** Cystathionine beta-synthase, s-adenosyl-L-methionine, allosteric regulation, activation, enzyme, filament, cryo electron microscopy, conformational change, aggregation, homocystinuria

## Abstract

Cystathionine beta-synthase (CBS) is an essential metabolic enzyme across all domains of life involved in the production of glutathione, cysteine, and hydrogen sulphide^1–4^. Human CBS appends to its conserved catalytic domain a regulatory domain that modulates activity by S-adenosyl-L-methionine (SAM) and promotes oligomerization^5–12^, however the molecular basis is unknown. Here we show using cryo-electron microscopy that full-length human CBS in the basal and SAM-bound activated states polymerises as filaments mediated by a conserved regulatory domain loop. In the basal state, CBS regulatory domains sterically block the catalytic domain active site, resulting in a low activity filament with three CBS dimers per turn. This steric block is removed when in the activated state, one molecule of SAM binds to the regulatory domain, forming a high activity filament with two CBS dimers per turn. These large conformational changes result in a central filament of SAM stabilised regulatory domains at the core, decorated with highly flexible catalytic domains. Polymerization stabilises CBS and increases the cooperativity of allosteric activation by SAM. Together our findings elaborate our understanding of CBS enzyme regulation, and open new avenues for investigating the pathogenic mechanism and therapeutic opportunities for CBS-associated disorders^3,13–17^.

---

The pyridoxal 5’-phosphate (PLP)-dependent enzyme CBS^18^ catalyses the condensation of serine and homocysteine to form cystathionine. Playing a pivotal role in the transsulfuration pathway and redox regulation^18^ and being linked to one carbon metabolism, CBS produces cysteine, glutathione as well as the gaseous transmitter hydrogen sulphide^1^ (H_2_S). CBS has recently gained interest as a therapeutic target as inhibition of H_2_S production has been suggested for treatment of specific cancers^16^ and Down’s Syndrome^15^. Inherited lost-of-function mutations of CBS result in classical homocystinuria (HCU), the most common inborn error of sulphur metabolism^17^. Classical homocystinuria has been recognised as a protein misfolding disorder as many of the known pathogenic mutations result in aggregation and degradation of the CBS enzyme^19,20^.

Human CBS adopts a unique multi-domain structure: an N-terminal heme binding domain (residues 38-74), a central PLP-dependent catalytic domain (CD, residues 75-382), and a C-terminal regulatory domain (RD, residues 411-551)^2,5,10^ (Fig. 1a). The RD adopts the evolutionarily conserved Bateman module, consisting of two tandem *CBS* motifs (*CBS-1* and *CBS-2*) and a surface-exposed loop (residues 516-525) extending from a *CBS-2* motif (Extended Data Fig. 1a). Bateman modules act as a regulatory sensor in response to the binding of predominantly adenosine-containing ligands^21^, and are present in enzymes across the three domains of life^22^. In mammals, the activity of CBS is increased by the binding of SAM to the Bateman module^23^. The enzyme is therefore proposed to transition between the basal state in the absence of SAM, and the SAM-bound activated state^23^

**Fig. 1.**
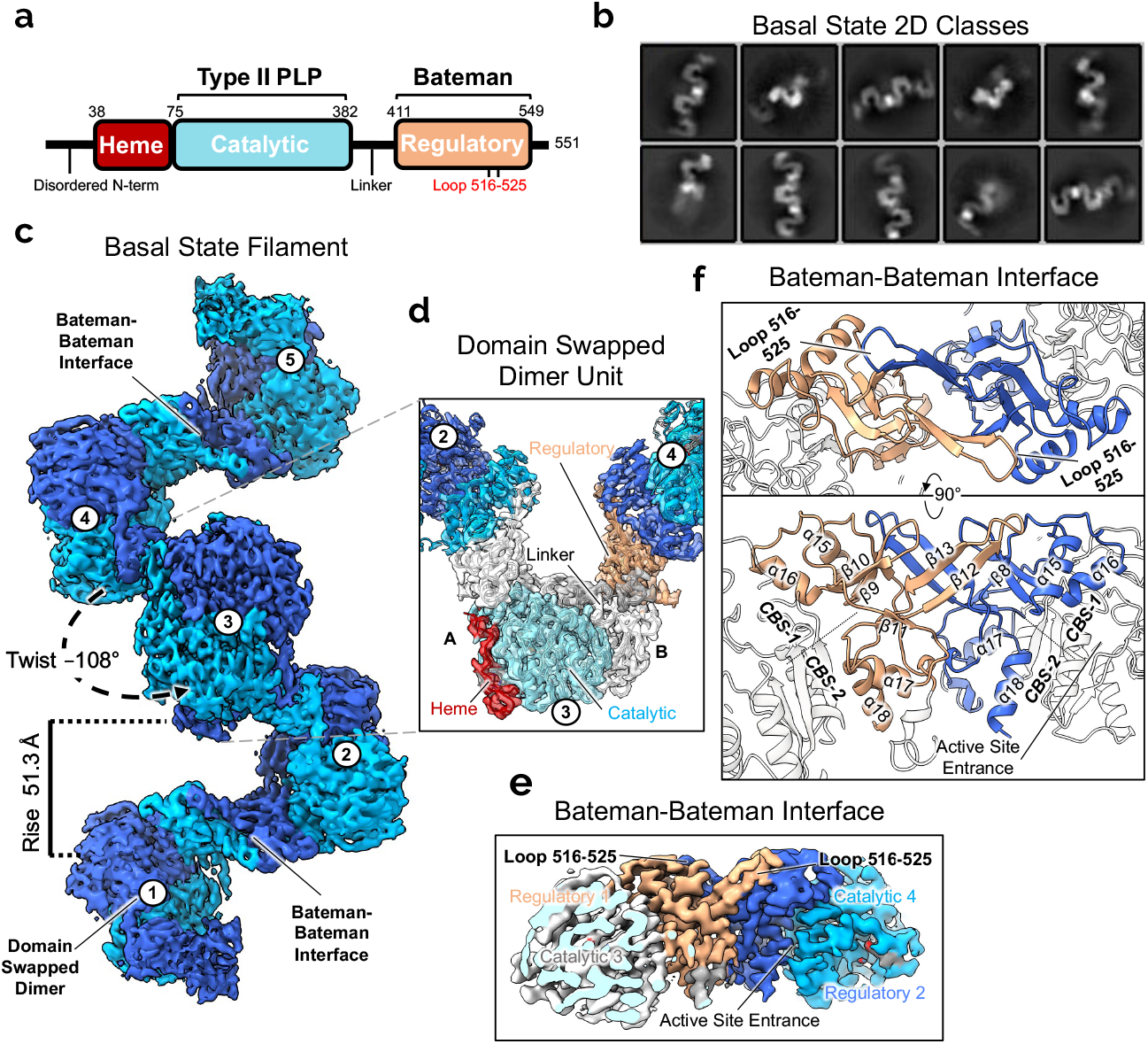
Cryo-EM structure of the basal state of CBS. **a**, Domain diagram of human CBS. **b**, Initial representative 2D classes of CBS in the basal state. **c,** Cryo-EM helical reconstruction of the CBS filament in the basal state at a resolution of 3.9 Å. Individual domain swapped dimers are numbered. **d,** Single particle reconstruction focused on one CBS dimer unit at a resolution of 3.0 Å. One monomer is coloured as in Fig. 1a. **e**, Close-up view of the EM density at the filament interface. This is formed by regulatory domain interactions from neighboring domain-swapped CBS dimers. The active site entrance of one catalytic domain is indicated. **f**, Model of the Bateman-Bateman interface formed by regulatory domain interactions. The C2 symmetry is apparent and shows the key role of the loop 516-525. The active site entrance of one catalytic domain is indicated.

Crystal structures of human CBS^10,8,9^ have been determined of CD alone (CBS^CD^), of full-length protein engineered with the RD loop deletion (CBS^Δ516–525^)^8,12^, and of RD alone with the loop deletion (CBS^RDΔ516–525^) complexed with SAM. Together, they depicted human CBS as a homodimer, where both CD and RD can dimerise independently. The structural data also revealed the molecular mechanism of SAM activation, whereby the Bateman module in the RD acts as an autoinhibitory cap to the CD, and upon binding SAM undergoes a conformational change relieving the inhibition to allow active site access for catalysis (Extended Data Fig. 1)^9,10^. However, these structures do not take into account various reports that the RD promotes the formation of CBS tetramers and higher order oligomers with a high tendency to aggregate^7,24–26^. Indeed, CBS oligomerization and SAM response varies across the animal kingdom^6^; it is unclear how human CBS oligomerises and its relation to SAM allosteric regulation is unknown^21,22^.

Additionally, mutations on or deletion of the Bateman module can rescue the most common homocystinuria associated mutations, but the structural information so far has given limited insight^27–29^. Efforts have also been made on developing new small molecule therapies targeting CBS^3,14,15^, but the lack of full-length enzyme structure has been a possible hinderance. Here we used a combination of biophysical techniques and cryo-electron microscopy showing that human CBS full-length protein polymerises as a filament adopting two very different morphologies dependent on SAM binding, and that polymerization plays a role in both enzyme activation and stability.

## Cryo-EM of full-length human CBS reveals a filamentous architecture

We pursued the cryo-electron microscopy structure of full-length CBS initially using two constructs: one where an N-terminal His-tag was removed during purification (CBS^FL^), and one with a permanent C-terminal His-tag (CBS^FL-CHis^) (Extended Data Fig. 2a). In analytical gel filtration on a Superose 6 column, both CBS^FL^ and CBS^FL-CHis^ proteins eluted as a broad peak, suggestive of different oligomeric states with molecular weights larger than tetramers (Extended Data Fig. 2b). Similar observations of larger-than-tetrameric CBS^FL^ oligomers, and even higher molecular weight oligomers for CBS^FL-CHis^, were made in Coomassie-stained clear-native and blue-native PAGE experiments (Extended Fig. 2c, d). As control, CBS^Δ516–525^, CBS^CD^, and CBS^RDΔ516–525^ behaved as expected dimers^30^. Micrographs of full-length CBS from both constructs in vitreous ice clearly showed the presence of flexible filaments (Supplementary Fig. 1a, 3a) of varying lengths agreeing with 2D classes (Fig. 1b, Supplementary Fig. 1b, 3b). Multiple maps were processed using both helical (Supplementary Fig. 1, 3) and single particle (Supplementary Fig. 2, 4) reconstruction to resolutions of between 3.9 to 3.0 Å. Both constructs resulted in maps with virtually identical conformations (Extended Data Fig. 3a), and both were used for modelling.

The CBS filament adopts a left-handed helical architecture with a twist of −108° and rise of 51 Å (Fig. 1c, Supplementary Fig. 1, 3). The domain-swapped dimer, previously observed in crystal structures, is the repeating unit and helical formation is driven by inter-ic RD interactions, i.e., between the RDs of neighbouring s of the filament. Here the RD sits atop the active site entrance of the catalytic domain hindering substrate access (Fig. 1c, e). Loop 516-525, a β-turn-β extension from the *CBS-2* motif in the conserved Bateman module, plays a key role in driving oligomerization. Specifically, one RD each from two neighbouring CBS s interact in a butterfly-like dimeric arrangement, related by C2 symmetry, such that the loop 516-525 from a RD of one dimer forms a clasp around a RD of the neighbouring dimer. This packing arrangement, burying a total surface area of ~1550 Å^2^ (Fig. 1e, f, Extended Data Fig. 3b), is stabilised through two interfaces composed entirely of the RDs. The first interface is between the *CBS-2* and *CBS-1* motifs, involving loop 516-525 of one dimeric subunit and α-helix 15 of the neighbouring dimeric subunit. Here, main-chain interactions are mediated between β-strand 12 residues 516-519 of one RD and β-strand 8 residues 422-426 of the adjacent RD (Fig. 1f, Extended Data Fig. 3d). This arrangement results in residue Tyr518 from the oligomerization loop slotting into a hydrophobic pocket formed by Leu423, Val425, Ile429, and Ile437 of α-helix 15 (Extended Data Fig. 3d). The second interface involves inter-ic RD contacts, between the two neighbouring *CBS-2* motifs. Here RD residues Leu419, Leu492, Met529, and Phe531 of one dimeric subunit form hydrophobic packing interactions with the equivalent residues of the neighbouring dimeric subunit (Extended Data Fig. 3e). Due to these interactions both Ala421 and Pro422 are shifted from their positions as observed in the previous crystal structures of CBS^Δ516–525^ (Extended Data Fig. 3f)^8–10^. Altogether, the two modes of inter-dimeric RD interactions demonstrate how the loop 516-525 is the main driver of CBS oligomerization^10,13,20,29^.

## CBS degrades into tetramers and dimers

Though both full-length CBS constructs resulted in highly similar filament maps, one key difference was the observed protein degradation and the presence of slightly shorter oligomers of the CBS^FL^ construct without a permanent His-tag (Extended Data Fig. 2a, b). Consequently, along with the filament classes we observed 2D averages of this construct that represented these degradation products (Extended Data Fig. 4a, b). One collection of classes appeared to be the catalytic domain dimer alone, suggesting that the RD was degraded from some full-length protein, although we could not obtain a reasonable reconstruction due to its small size (~80 kDa) (Extended Data Fig. 4a). Another collection of classes resulted in a 3.8 Å map of a degraded CBS tetramer (Extended Data Fig. 4a, b, Supplementary Fig.4). Here two degraded heterodimers, composed of one full-length protomer and one RD-degraded catalytic domain protomer (Extended Data Fig. 4b), interact through a single Bateman-Bateman interface, highly similar to the arrangement seen in the intact filament. The degradation of the RD renders the active site loops within the catalytic domain of the full-length CBS in a more opened state (Extended Data Fig. 4c).

## SAM binding transforms the morphology of the CBS filament

The global methyl donor SAM functions as an allosteric activator of human CBS activity^23^. In our biophysical characterisation assays (Extended Fig. 2c, d), SAM had no significant effect on the oligomeric status of all CBS constructs, agreeing with previous reports^12,31^. Since oligomerization occurs without SAM, we hypothesised that SAM could modulate the morphology of the CBS filament, as part of its role as an allosteric activator. To this end, cryo-EM was used to analyse CBS^FL-CHis^ in the presence of SAM. Micrographs showed that CBS retains a filamentous architecture in the presence of SAM, but with a significantly altered morphology as shown by 2D classes (Fig. 2a, Supplementary Fig. 5a, c). Obtaining a 3D reconstruction took considerable effort due to the highly flexible nature of the filaments (Supplementary Fig. 5b). Through helical refinement we generated a global map at 4.0 Å resolution which reveals a central helical stalk decorated with highly flexible lobes (Fig 2b). To aid in model building, we applied masks and performed local refinement that resulted in local maps for the central stalk at 4.1 Å resolution and for the flexible domain at 8.3 Å resolution (Supplementary Fig. 6).

**Fig. 2.**
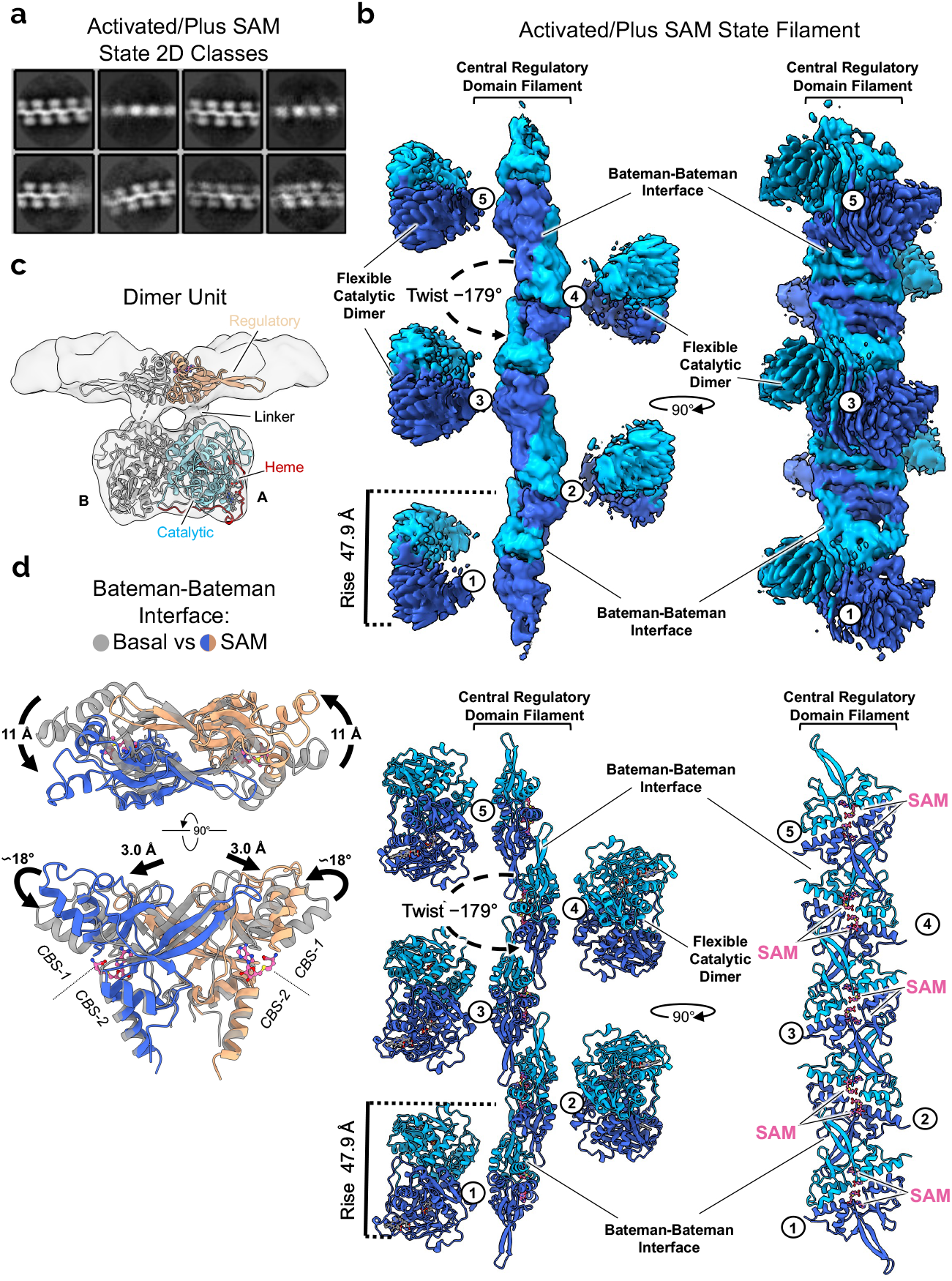
Cryo-EM structure of the SAM bound activated state CBS. **a,** Initial representative 2D classes of the activated CBS filament in the presence of SAM. **b**, Global helical cryo-EM map of the activated SAM bound CBS^FL^ filament at a resolution of 4.1 Å and atomic model. The catalytic domains are omitted for clarity in the bottom right panel. Individual dimers are numbered. **c**, Single particle reconstruction focused on one CBS dimer unit at a resolution of 8.0 Å. One monomer is coloured as in Fig. 1a. **d**, Structural alignment of the Bateman-Bateman interface in the basal and activated states showing the conformational change due to SAM binding.

These maps allowed us to model the entire filament, by docking one catalytic domain dimer into each flexible lobe (PDB: 4PCU), and repeating units of the regulatory domain dimer (PDB: 4UUU) into the central stalk. The overall morphology of the resulting filament is drastically different in comparison to the basal state, as reflected by the 66% increase in twist (−178.6°) and 10% decrease in rise (46.9 Å) of the filament in the presence of SAM (Fig. 2b, Extended Data Fig. 5a). Previous crystal structures of the non-filamentous CBS^Δ516–525^ dimers^9,10^ showed that SAM binding to the RD elicits its dis-association from the CD and subsequently its homo-dimerization, thereby freeing the active site access for catalysis (Extended Data Fig. 1a). In the context of the full-length enzyme elucidated here, the SAM-mediated conformational change creates a central filament stalk composed of repeating units of the SAM-bound RD dimer, arranged in an antiparallel “daisy-chain” like fashion due to the interactions of the loop 516-525 with the neighbouring subunit (Fig. 2b, c). The central filament stalk is decorated by highly flexible CDs, where the active site entrance loops are free to open and hence increase the accessibility for substrates (Extended Data Fig. 5b)^9^.

Comparing our basal and activated filaments, the SAM-mediated conformational change to the RD is highly agreeable with that observed in the crystal structures of CBS^Δ516–525^ dimers where there is a relative 18° rotation between the *CBS-1* and *CBS-2* motifs caused by SAM binding (Fig. 2d)^9,10^. Observing this interface of *CBS-1* and *CBS-2* in isolation, the conformational change is mainly localised in the *CBS-1* motif and appears as an unfurling of the “butterfly wings” of this dimeric arrangement (Movie 1). This results in α-helix 15 displaced by 8 Å and α-helix 16 by as much as 11 Å from their original positions. Interestingly loop 516-525 from the neighbouring subunit also moves 3.0 Å to maintain its interactions with residues 422-426 and α-helix 15 (Fig. 2d). Additionally, alignment to the central helical Z-axis and a simple morph of the global basal and activated models show that the large conformational change is possible in the filament with little clashes when the CD moves in concert with the RDs (Extended Data Fig. 5d, Movie 2)^9,10^.

## Filament formation does not create more SAM binding sites in CBS

Each Bateman module, assembled from the tandem *CBS-1* and *CBS-2* motifs, contains in principle two ligand-binding sites (S1 and S2) related by dyad symmetry^8^. In our activated filament maps, ligand density was present for SAM only at the S2 site (Fig. 2b, 3a). No apparent density was found for SAM at the S1 site nor at the filament interfaces. This 1:1 (one SAM to one CBS protomer) stoichiometry is identical to previously observed CBS^Δ516–525^ crystal structures where only the S2 site was occupied by SAM^9,10^. These structures suggest that Phe443 and Asp538 at the S2 site are key residues in SAM binding^10^. Therefore, to confirm the 1:1 binding of SAM we purified the mutants F443A and D538A, and initially tested their enzymatic response to SAM. These mutants showed a low basal activity like wild-type, but unlike wildtype could not be stimulated by SAM (Fig. 3b).

**Fig. 3.**
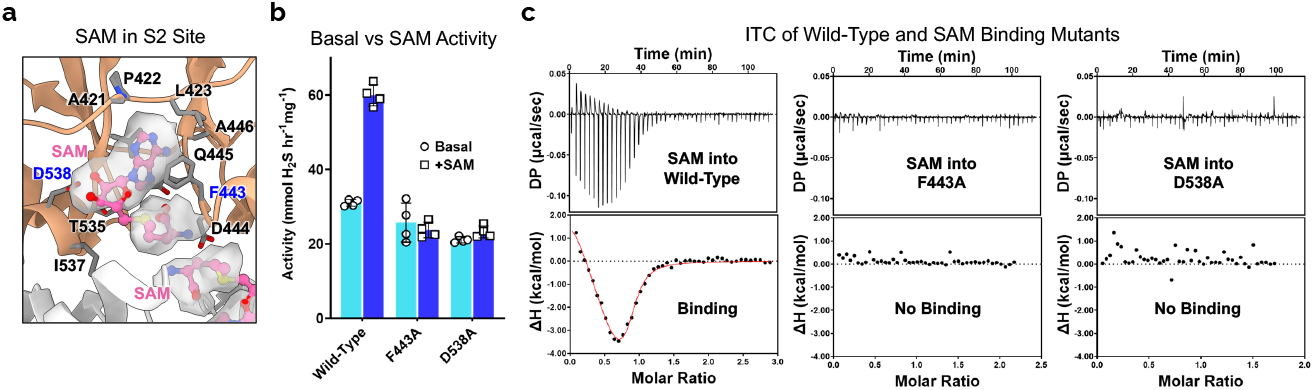
Full-length CBS binds only one SAM per monomer. **A,** Cryo-EM density of SAM bound in the S2 site. Interacting residues are represented as sticks and coloured gray. SAM is represented as balls and sticks and is coloured pink. **b**, H_2_S producing activity of wild-type, F443A, and D538A CBS^FL-CHis^ as purified (basal) and in the presence of 300 μM SAM (+SAM). Mean values and error bars are ± s.d. of *n* = 4 technical repeats. **c**, ITC titrations of SAM against wild-type, F443A, and D538A CBS^FL-CHis^. Each titration is representative of two independent experiments.

We next performed isothermal titration calorimetry (ITC) of wild-type CBS^FL-CHis^ against SAM which demonstrated two apparent binding events at ~160 nM and ~600 nM, in agreement with previous reports^11,12,31^. In contrast the S2 site mutations, F443A and D538A, eliminated both SAM binding events (Fig. 3c, Extended Data Fig. 6b, d), again suggesting that no other SAM sites exist in the filament. We reasoned that the two apparent binding events are attributed to the binding of one SAM to the S2 site and the resulting conformational rearrangement into the activated state. In support of this interpretation, ITC of CBS^FL^ and CBS^Δ516–525^ (where the protein can respond to both ligand binding and conformational changes) both demonstrated two apparent binding events, whereas the RD alone protein CBS^RDΔ516–525^ (which only responds to ligand binding) presented only one event at ~4 μM (Extended Data Fig. 6c, d). Overall, our data indicates that CBS binds SAM in a 1:1 manner, and that filament formation does not create additional SAM binding sites.

## Filament formation increases cooperativity of SAM activation and CBS stability

Polymerization of metabolic enzymes, such as involving filament formation, has been linked to their regulation and stabilization^32,33^, and we hypothesised that the assembly of human CBS into a filament could play a similar role that facilitates enzyme catalysis. Our observation of loop 516-525 moving in tandem with the conformational change of α-helix 15 (Fig. 2d), in the presence of SAM, suggests potential crosstalk communication between regulatory domains from neighbouring CBS proteins (inter-dimer) in the filament (Movie 1). To investigate potential cooperativity within the CBS filament, we characterised CBS activity by measuring H_2_S production from the condensation of cysteine and homocysteine. Km values for both homocysteine and cysteine were essentially identical for the four constructs at ~0.3 and ~20 mM respectively. CBS^FL^, CBS^FL-CHis^, and CBS^Δ516–525^ were allosterically activated by SAM whereas CBS^CD^ was not (Extended Data Fig 6a). By titrating SAM, we found that the responsive constructs showed a ~2-fold increase in activity, exhibiting Kact for SAM of ~26.0-36.0 μM (Fig. 3a, Extended Data Fig 6b) agreeing with reported values^11,25,34^. The activation of CBS^FL^ and CBS^FL-CHis^ was cooperative with a Hill coefficient (*n*_Hill_) of 3.0-3.6, whereas the non-filamentous CBS^Δ516–525^ presented a lower *n*_Hill_ of 2.0 (Fig. 3a, Extended Data Fig 6b).

Next, we determined if filament formation alters CBS stability by using thermal shift. The CBS^Δ516–525^ protein, which does not form filaments, is less thermostable than CBS^FL^ by ~5 °C, confirming that filamentation increases stability. We also found that CBS^FL-CHis^ is more thermostable than CBS^FL^ by ~3 °C (Fig. 3b, Extended Data Fig. 5a), which may be due in part to it forming longer oligomers and being less degraded than CBS^FL^ (Extended Data Fig. 2)^30^. Moreover, it is known that the activity of human CBS can be increased by thermal activation, likely due to the denaturation of the regulatory domain that relieves its autoinhibitory effect^7,11,23^. Repeating this assay, on both CBS^FL^ and CBS^Δ516–525^ we determined Tm values of 49.5 °C and 43.7 °C respectively showing that CBS^Δ516–525^ is more prone to thermal activation and that its regulatory domain is less stable (Fig. 3c). These findings suggest that polymerization alters both the cooperativity of SAM activation and stabilizes the regulatory domain probably to maintain CBS in the basal state conformation.

## Discussion

Conflicting reports of human CBS oligomerization, evidenced by a variety of biophysical and immunoblotting techniques, using both recombinant and endogenous sources of the protein, have plagued the literature since its initial characterisation^5,12,20,24–26,35,36^. Here we have shown that human CBS oligomerises into filaments, adopting (at least) two distinct architectures respectively for the basal and activated states. Chiefly our findings reflect the initial reports of CBS purified from human liver, where it was shown to form large oligomers with a tendency to both aggregate and degrade to a more active state^26^. This discovery of filamentation evaded past crystallographic studies involving an engineered protein that removes a surface loop (residues 516-525)^8–10^ now revealed to be key to polymerization, alongside the assumption that CBS is predominantly a tetrameric protein^6^. Our observation of CBS filaments with heterogenous lengths in cryo-EM micrographs therefore sufficiently explains previous findings of a mixture of CBS oligomers. Due to this heterogeneity the reported tetrameric state of CBS is likely to be two dimers interacting through a single Bateman module pair but additionally could also be formed via the degradation of the Bateman module that we have shown here (Extended Data Fig. 4).

The link between the oligomeric state and SAM activation has also had conflicting reports. Originally shown to bind only one SAM molecule per monomer, more recent ITC studies suggested a two-site model where a kinetically stabilizing high-affinity SAM binding site would be formed from oligomerization, while enzyme activation is driven by SAM binding to the lower affinity S2 site observed in dimeric CBS^Δ516–525^ crystal structures^11,12,37^. Our cryo-EM structure of SAM-bound activated state (Fig. 2), mutagenesis data (Fig. 3), and previous biophysical analyses^10^ all conform to the notion that only the S2 site exists to bind SAM. Though our ITC analysis of full-length CBS fits the two-site model (Fig. 3c, Extended Data Fig. 6), we reason that the thermograph reflects not only SAM binding but also the structural rearrangements that have to occur for activation. This transition from basal to activated states requires a significant conformational change and likely follows a multistep process (Extended Data Fig. 5d, Movie 2). Conformational changes due to ligand binding are known to be a major contributor to heat capacity changes^38^ and there is precedence for ligand binding to Bateman modules to diverge from a simple one-site binding model when dimerization of Bateman modules occurs^39^. Therefore, we regard our data as relative measurements of both SAM binding and conformational changes. As the kinetically stabilising site has been suggested for drug discovery^11,37^, our findings here suggest that alternative frameworks in the context of filament should be considered for targeting the CBS regulatory domain (discussed below).

This study now firmly places human CBS in the growing membership of filamentous metabolic enzymes (Fig. 5a). Higher order oligomerization in response to signal transduction from ligand (nutrient) binding or stress has been shown for many eukaryotic metabolic enzymes such as acetyl-CoA carboxylase (ACC), inosine-5-monophosphate dehydrogenase (IMPDH), and cytidine triphosphate synthase (CTPS)^32,33^. The yeast CBS orthologue, *S. cerevisiae* Cys4p, forms punctate foci during stationary phase in response to nutrient levels^40^, an observation that suggests filamentation^33^. However yeast Cys4p, contrary to human CBS, does not undergo allosteric feedback activation in response to SAM^6^ and does not contain the same oligomerization loop in the regulatory domain (Extended Data Fig 8). Therefore, it remains unclear if SAM constitutes the signal or driver for Cys4p oligomerization. For human CBS, however, filament formation takes place both in the absence and presence of SAM. AlphaFold predictions^41^ and sequence alignment of Bateman modules from various CBS orthologues suggests that filament formation is possibly conserved in chordates (Extended Data Fig. 8, 9). This therefore implies that CBS oligomerization could be an evolutionarily strategy overall but could involve different structural components and serve different functional outcomes.

For many metabolic enzymes, filamentation and oligomerization serve to provide new ligand binding sites, generate unique catalytic conformations, or transduce signals across multiple enzyme subunits. Such functional modification to the enzyme is often reflected in a significant increase (or decrease) in cooperativity and activity upon filament formation^32,33^. For the case of human CBS, we observe only a modest increase in cooperativity in comparison to the loop deleted CBS^Δ516–525^ construct (Fig. 4a). This is not surprising as the CBS filament interface involves a single point of contact (Fig. 1, 2). In other enzymes, such as IMPDH which forms a filament of tetramers, multiple contacts are formed across single filament interfaces with corresponding high values of cooperativity^42^. We do however find that the full-length CBS filament is more stable and less prone to thermal activation than the loop-deleted CBS^Δ516–525^ dimer (Fig. 4b, c). We previously observed that the CBS^Δ516–525^ construct is conformationally flexible without SAM, presenting as two populations in ion mobility experiments^10^. Thus, we propose that the primary objective of filament formation in CBS is to increase the kinetic stability of the regulatory domain to maintain the basal conformation of the enzyme^11,31,37^ (Fig. 5a). The notion of filament formation increasing stability is further supported by the His-tagged construct that presents longer polymers, higher stability, and less degradation than the construct without a His-tag (Fig. 4b, Extended Data Fig. 2). Increased stability and activity of human CBS due to a C-terminal His-tag has been previously reported^30^. Unfortunately, we cannot structurally rationalize this effect as we observed no interpretable density in the maps of this construct for the affinity tag at the Bateman-Bateman interface (Fig. 1, 2). We theorise though that the His-tag could be acting as a proxy for an as yet unidentified ligand that may regulate CBS oligomerization. Further investigations are clearly warranted; however, these observations show that filament formation can be modulated (positively or negatively) by changes at the Bateman-Bateman interface.

**Fig. 4.**
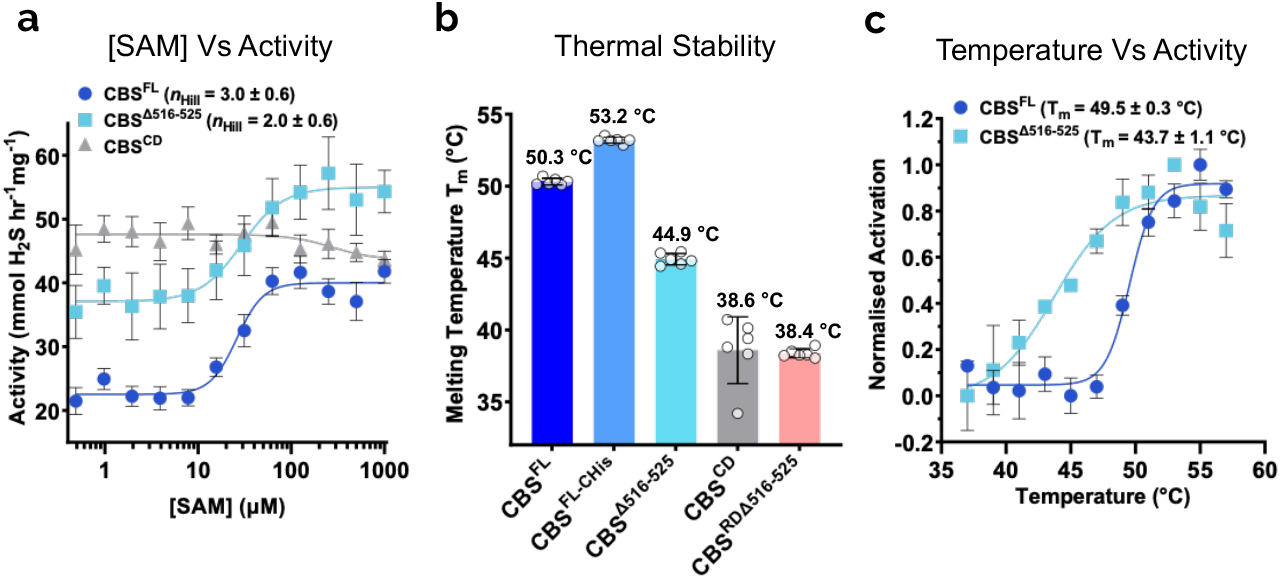
Filament formation increases SAM cooperativity and stability of CBS. **a**, H_2_S producing activity of CBS^FL^, CBS^Δ516–525^, and CBS^CD^ in response to increasing amounts of SAM. Mean values and error bars are ± s.d. of *n =* at least 4 technical repeats. **b**, Thermal stability of CBS^FL^, CBS^FL-CHis^, CBS^Δ516–525^, CBS^CD^, and CBS^RDΔ516–525^. Mean values and error bars are ± s.d. of *n* = 6 technical repeats. **c**, Thermal activation of CBS^FL^ and CBS^Δ516–525^ activity. Mean values and error bars are ± s.d. of *n* = 4 technical repeats.

**Fig. 5.**
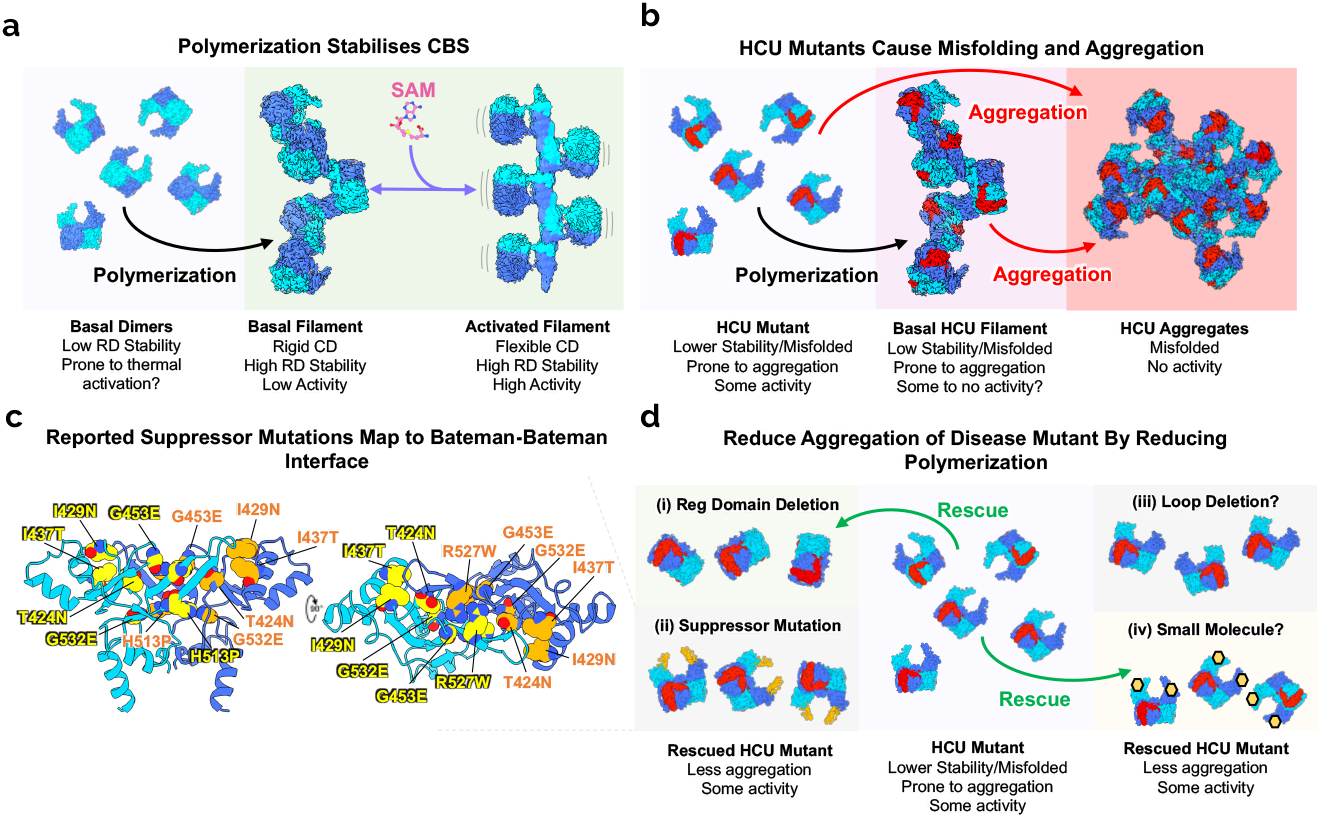
Filament formation stabilizes CBS but potentially increases the aggregation of disease mutants. **a,** Proposed model of CBS stabilization by polymerization and allosteric activation. **b**, Proposed model of HCU mutant CBS polymerization and its relation to CBS aggregation. Red indicates mutation on the catalytic domain. **c**, Previously reported HCU mutant suppressor mutations are located at the Bateman-Bateman interface. **d**, Proposed model of HCU mutant rescue by reducing polymerization and aggregation. Rescue has been shown by (i) deletion or (ii) mutation of the catalytic domain, which may sacrificially reduce CBS’s natural propensity to polymerize into filaments. The (iii) deletion of the loop 515-526 or the (iv) binding of a small molecule at the Bateman-Bateman interface could also reduce polymerization and reduce aggregation of HCU mutants.

Inherited mutations in CBS result in classical homocystinuria (HCU), in which most recorded mutations are missense^4^, and the dominant molecular mechanisms have been recognised as protein misfolding and aggregation^17,19,20^ (Fig. 5b). Rescue of mutant CBS activity has been documented by chemical chaperones^35,43^, heme arginate^44^, and proteostasis inhibitors^45^, suggesting a small molecule therapy could be developed^46,47^. It is intriguing that genetic suppression in a yeast model of the disease has also been reported, where deletion^29^ or missense mutations on the regulatory domain^29^ can overcome the deleterious effects of the most common HCU mutations. Disease associated mutants in general can produce hydrophobic patches in the protein due to local misfolding, that will result in aggregation^48–50^. It is probable that many HCU mutations generate hydrophobic patches on the catalytic domain resulting in further non-specific interactions that lead to aggregation^19^ (Fig. 5b).

Considering our findings, we hypothesise that a mechanism of rescue could be the reduction of the natural propensity for CBS to polymerise which could reduce one pathway towards aggregation (Fig. 5d). In support of our reasoning, 1) deletion of the regulatory domain prevents filament formation (Extended Data Fig. 2), and 2) the seven reported suppressor mutations from the yeast model of disease can be all mapped onto the hydrophobic face of the regulatory domain that forms the Bateman-Bateman interface (Fig. 5c, Extended Data Fig. 10a). All seven residues are conserved in chordate CBS (Extended Data Fig. 8) and are predicted to alter interactions at the oligomeric interface (Extended Data Fig 10b). The original report suggested that these mutants altered the interaction between the regulatory and catalytic domains trapping CBS in a partially open conformation which is non-responsive to SAM. However, in the background of the wild-type enzyme no slight increase in basal activity was observed^28^ and as such we believe that altered oligomerization should be considered as an aspect of rescue. As dimeric CBS^Δ516–525^ behaves almost like full-length (i.e., is active and SAM responsive), a small molecule that disrupts CBS polymerization and reduces aggregation could be a potential therapeutic avenue for the treatment of HCU (Fig 5d).

Overall, we have determined multiple structures of human CBS showing that the fulllength enzyme polymerises as an active filament which changes conformation due to SAM. Future work should consider further the role of CBS polymerization in protein misfolding and aggregation. It is interesting that there are catalytic domain HCU mutations that result in a CBS enzyme with normal basal activity but non-responsive to SAM^43^. SAM non-responsive HCU mutations in the regulatory domain exhibited an enzymatic activity closer to the activated state^34^, suggesting that some mutants may lock CBS in one conformation. Cryo-EM studies of these and other disease associated mutants will give insight into the molecular mechanism of protein misfolding of CBS and may have implications in understanding the misfolding of other multidomain metabolic enzymes.

## Methods

### Cloning, expression, and purification of human CBS proteins

The gene for human CBS (UniProt P35520) was cloned into pNIC-Bsa4 and pNIC-CH encoding for a TEV cleavable N-terminal and permanent C-terminal His-tag respectively. The constructs CBS^Δ516–525^ and CBS^CD^ (residues 1-413) along with single point mutations of CBS were introduced using In-Fusion (Takara) or QuikChange (Agilent) mutagenesis and confirmed by sequencing. CBS was expressed in *E. coli* Rosetta (DE3) cells in auto induction Terrific Broth (TB) supplemented with 50 μg/ml kanamycin, 34 μg/ml chloroamphenicol, 0.3 mM δ-aminolevulinic acid, 0.0025% pyridoxine–HCl, 0.001% thiamine–HCl, and 0.1 mM ferric chloride at 30 °C, 200 rpm for 24 hours. Cells were resuspended in lysis buffer (50 mM sodium phosphate, pH 7.5, 500 mM NaCl, 0.5 mM TCEP, 5% glycerol, 1.0% Triton X-100, 0.1 mM PLP, 2 mg/ml lysozyme) and lysed by sonication. CBS proteins with a TEV cleavable N-terminal His-tag were purified using Ni-NTA agarose (Qiagen) resin and were treated to gel filtration using a Superose 6 Increase 16/600 column or Superdex 200 Hiload 16/600 column (Cytiva) equilibrated in storage buffer (25 mM HEPES, pH 7.5, 500 mM NaCl, 0.5 mM TCEP, 5% glycerol). Fractions containing CBS protein were pooled and treated with His-tagged TEV protease overnight at 4 °C, and then passed over Ni-NTA agarose resin to remove the TEV protease and uncleaved protein. CBS proteins with a permanent C-terminal His-tag were purified using TALON (Clontech) resin followed by anion exchange using a Hitrap Q column (Cytiva). Anion exchange elutions were polished by gel filtration using a Superose 6 Prep Hiload 16/600 column (Cytiva) equilibrated in storage buffer (25 mM HEPES, pH 7.5, 500 mM NaCl, 0.5 mM TCEP, 5% glycerol). For all purifications appropriate fractions were pooled, concentrated to 5-20 mg/ml, snap frozen, and stored at −80°C.

### Clear and blue native-PAGE

Clear and blue native-PAGE was carried out according to manufacturer’s instructions (Life Technologies). CBS constructs were diluted in 25 mM HEPES, pH 7.5, 200 mM NaCl, 2.0 mM TCEP. In some cases, 1 mM SAM was added and incubated at room temperature for 5 minutes before loading. All samples were at 1.0 mg/ml and 8.0 μg total protein.

### Analytical SEC

Analytical SEC was carried out using a 10/300 GL Superose 6 Increase column (Cytiva) equilibrated in 25 mM HEPES, pH 7.5, 500 mM NaCl, 0.5 mM TCEP, 5% glycerol. 250 μl of each CBS construct was loaded at 3.0 mg/ml with a flow rate of 0.3 ml/min. 500 μl fractions were collected for analysis by SDS-PAGE.

### Grid preparation and cryo-electron microscopy

CBS^FL^ and CBS^FL-CHis^ were diluted to 1.0 mg/ml (15 μM) into 25 mM HEPES, pH 7.5, 200 mM NaCl, 2.0 mM TCEP, 0.005% (v/v) tween-20 for the basal state. For the activated state CBS^FL-CHis^ was diluted to 0.75 mg/ml (11.25 μM) into 25 mM HEPES, pH 7.5, 200 mM NaCl, 2.0 mM TCEP, 0.005% (v/v) tween-20, 5 mM SAM. Grids were prepared using a FEI Vitrobot Mark III (Thermo Fisher Scientific) at 4 °C and 100% humidity. 3 μl of sample was applied to a plasma treated gold coated R 1.2/1.3 300 mesh holey carbon grid (Quantifoil), with a blot force of 0, a blot time of 3 seconds and a wait time of 10 seconds.

Movies of the CBS^FL-CHis^ basal state were collected at eBIC (Diamond Light Source) on a Titan Krios equipped with a Falcon 3EC direct electron detector (Thermo Fisher Scientific) operating in counting mode. Images were imaged at 300 kV with a magnification of 75,000×, corresponding to a pixel size of 1.085 Å. 40 frames over 60 seconds were recorded with a defocus range of −0.9 μm to −3.0 μm with a total dose of 37.85 e^−^ A^−2^ (0.823 e^−^ A^−2^ per frame). A total of 1,740 movies were collected in a single session. MotionCor2^51^ was used to correct beam induced motion and CTF was estimated using CTFFIND-4.1^52^. For helical reconstruction, particles were picked using the filament picker of Relion 3.0.8^53^ resulting in 239,739 particles extracted. Helical picks were subjected to multiple rounds of 2D classification producing 82,810 particles that were imported to CryoSPARC-3.1.0^54^. Further 2D classification to remove any junk particles reduced this to 76,663 particles. Helical parameters were roughly determined from a low resolution Glacios collected map. Nonuniform helical refinement with D1 symmetry (C2 symmetry perpendicular to the helical axis) imposed resulted in 3.7 Å map with refined helical twist of −108.4 ° and rise of 51.2 Å. For single particle analysis, particles were auto-picked using the Relion 3.0.8^53^ (Laplacian of Gaussian function) resulting in 760,869 particles extracted. One round of 3D classification with 4x binned images and a model from a subset of the data was used to remove bad particles and contamination. This resulted in 392,163 particles that were unbinned and subjected to per particle CTF refinement and Bayesian polishing. After another round of masked 3D classification, one class consisting of 207,509 particles was identified to have the highest level of structural detail. A further round of CTF refinement and Bayesian polishing followed by masked auto-refining was used to produce particles for cryoSPARC-3.1.0^54^. 2D classification followed by heterogonous refinement reduced the number of good particles to 188,230. Multiple rounds of non-uniform refinement, local CTF refinement and local non-uniform refinement with C2 symmetry imposed resulted in a 3.0 Å map.

EER formatted movies of the CBS^FL^ basal state were collected at the York Structural Biology Laboratory (YSBL) on a Glacios equipped with a Falcon 4 direct electron detector (Thermo Fisher Scientific). Images were imaged at 200 kV with a magnification of 150,000×, corresponding to a pixel size of 0.935 Å. Movies over 5.18 seconds were recorded with a defocus range of −1.4 μm to −2.0 μm with a total dose of 50 e^−^ A^−2^. A total of 2,628 movies were collected in a single session. All movies were imported into cryoSPARC-3.3.2^54^ where they subjected to patch CTF estimation and patch motion correction. For helical reconstruction, particles were picked using the filament tracer resulting in 749,213 particles extracted. Multiple rounds of 2D classification to remove any junk particles reduced this to 98,993 particles. Particle curation based off CTF fit further reduced this to 89,761 particles. Initial helical parameters were determined from the CBS^FL-CHis^ map. Non-uniform helical refinement with D1 symmetry imposed resulted in 3.9 Å map with refined helical twist of −108 ° and rise of 51 Å. For single particle analysis, 1,190,611 particles were picked and extracted using template-based picking. Rounds of 2D classification resulted in 160,400 particles that were subjected to ab-initio reconstruction and heterogenous refinement with three classes. Two classes were further separately processed using local non-uniform refinement with C2 symmetry imposed resulted in a 3.8 Å maps of both degraded and non-degraded CBS.

Movies of the CBS^FL-CHis^ activated state (SAM bound) were collected at eBIC (Diamond Light Source) on a Titan Krios (Thermo Fisher Scientific) equipped with a K3 (Gatan) direct electron detector operating in super-resolution mode. Images were imaged at 300 kV with a magnification of 81,000×, corresponding to a pixel size of 0.53 Å. 44 frames over 3.53 seconds were recorded with a defocus range of −0.9 μm to −3.0 μm with a total dose of (0.908 e^−^ A^−2^ per frame). 11,220 movies were collected in a single session. All movies were imported into cryoSPARC-3.1.0^54^ where they were motion corrected and the CTF estimated using patch motion correction (Fourier cropped to 1.06 Å) and patch CTF estimation respectively. Processing the activated state took considerable effort and initially filaments were picked using the filament tracer without templates on 2,790 micrographs. The resulting best classes from 2D classification were then used for another round of filament picking. Another round of 2D classification and picking the best classes were used to produce templates representative of the two dominant views with different filament widths. Two separate rounds of filament picking on the entire dataset resulted in 2,374,969 and 2,865,445 particles, which were eventually merged into a pool of 492,224 particles after many rounds of 2D classification and removal of duplicate particles. Helical parameters were roughly determined by visual inspection of a low resolution Glacios collected map. Rounds of non-uniform helical refinement with D1 symmetry imposed were used to re-centre particles and remove duplicates within 44 Å resulting in 425,260 particles. One further round of non-uniform helical refinement with D1 symmetry imposed resulted in a map at 4.0 Å resolution of the full filament with a refined helical twist of −178.6° and rise of 46.7 Å. Subsequently this map was used to make a soft mask (10 Å dilation with a soft padding of 50 Å) of the central filament region. Imposing this as a static mask during non-uniform helical refinement with D1 symmetry imposed resulted in a 4.1 Å resolution of the central regulatory domain with a helical twist of −177.7 ° and rise of 47.2 Å. To improve the resolution of the central regulatory domain the particles were then subjected to a masked local refinement with D1 symmetry applied and a less soft central regulatory domain mask (10 Å dilation with a soft padding of 20 Å). This resulted in a map with a resolution of 4.1 Å. Though distortions due to the flexibility of this central region were apparent at the edges of the map the three dimeric repeats at the centre of the map had improved features. To improve the quality and resolution of the highly flexible catalytic domains, masked local refinement with D1 symmetry imposed with a soft mask of the most central catalytic and regulatory domains (6 Å dilation with a soft padding of 20 Å) was used. Here the global helical map was filtered to 20 Å and alignments only considered a resolution of up to 9 Å resulting in map of 8.3 Å resolution.

### Model fitting, refinement, and validation

For the CBS^FL^ basal state structures, initially three CBS^Δ516–525^ structures (PDB: 4COO) were fitted using Molrep^55^ and the missing loop 513-527 was manually built in Coot^56^. Multiple copies of the CBS^FL^ model were docked into each map as appropriate using Phenix^57^. Rounds of refinement in Phenix^57^ were then used to refine the structure with manual adjustments in Coot^56^. For the activated state multiple copies of the isolated Bateman domain from our basal structure was docked using Phenix^57^ into the central locally refined map, and then flexibly fitted into a 10 Å filtered map using Namdinator^58^. This was followed by a second round of flexible fitting using the non-filtered map. The crystal structure of the Bateman loop-deleted dimer bound to SAM (PDB: 4UUU) was used as a guide to dock SAM into the appropriate density. This model and our basal state model were then used as references for refinement of the entire SAM bound model using Isolde^59^ and Phenix^57^. For the 8.3 Å map of a single catalytic domain the CBS^CD^ structure (PDB: 4PCU) was manually fitted into the density using Chimera^60^. These models were then assembled into a pseudo-atomic model of the global 4.1 Å activated map. All models were validated using Molprobity^61^.

### Enzyme activity assay

Kinetic parameters were determined by monitoring hydrogen sulphide (H2S) production using the fluorescent probe 7-azido-4-methylcoumarin (AzMC)^62^. Assays were performed in 25 mM HEPES, pH 7.5, 200 mM NaCl, 5 μM PLP, 10 mM glutathione, 10 μM AzMC, with 0.01% triton-x 100, in 384 well black plates, as a final assay volume of 50 μl. A final concentration of 100 nM of each CBS construct was used. For Michaelis-Menten kinetics cysteine was varied 0-40 mM with a constant 10 mM homocysteine and homocysteine varied 0-10 mM with a constant 40 mM cysteine. 300 μM SAM was added when appropriate. SAM titration assays were performed with a final concentration of 10 mM homocysteine and 40 mM cysteine, and SAM was added at a range of 0-1.0 mM. Thermal activation was carried out with CBS protein at 1 μM in 25 mM HEPES, pH 7.5, 200 mM NaCl, 50 μM PLP as 50 μl aliquots treated at different temperatures using a VeritiPro thermal cycler (Thermo Fisher Scientific) for two minutes. Treated samples were then put into ice before activity was assayed with 10 mM homocysteine and 40 mM cysteine. All plates were preincubated with enzyme for 10 minutes at 37 °C before the addition of cysteine. Plates were sealed with and spun at 900 x g for one minute before loading into the plate reader. H2S production was monitored by fluorescence at 450 nm (λ_ex_ = 365 nm) using a OmegaSTAR (BMG Biotech) at 37 °C. Each plate was read for one hour with a reading every one minute and raw rates were determined using MARS software (BMG Biotech). Activity readings were calibrated using a standard curve of known H2S concentrations using sodium hydrosulfide hydrate as a H2S source. Kinetic analyses were done in GraphPad Prism.

### Thermal shift assay

CBS constructs were diluted in thermal shift buffer (25 mM HEPES, pH 7.5, 200 mM NaCl, 2.0 mM TCEP) to 0.3 mg/ml with SYPRO-Orange (Invitrogen) diluted 1000X in a total volume of 20 μl. A QuantStudio 3 RT-PCR machine (Thermo Fisher Scientific) was used to measure melting temperatures.

### Isothermal titration calorimetry

Purified CBS proteins were buffer exchanged into 20 mM HEPES, pH 7.4 using Zeba spin columns (Thermo Fisher Scientific) at 4 °C. To prevent precipitation CBS^FL-CHis^ and its mutants were buffer exchanged into 20 mM HEPES, pH 7.4, 0.01% triton X-100 whereas CBS^RD^ was exchanged into 20 mM HEPES, pH 7.4, 200 mM NaCl. The appropriate buffer was then used to dissolve SAM from stocks to 500 μM. A MicroCal PEAQ-ITC machine with v1.21 control software for data collection (Malvern Panalytical) was used to perform ITC. CBS constructs were tested at 30-60 μM monomer in a 200 μl sample cell and were injected with 0.4 μl followed by 44 × 0.8 μl of SAM with 150 s spacing at 25 °C. Heats of dilution were determined by separate runs of SAM injected into buffer alone. Integrated heats were fit to using the Microcal PEAQ-ITC analysis software v1.30 (Malvern Panalytical) to obtain n, Kd, ΔH, and −TΔS.

### Structural analysis using AlphaFold multimer

CBS sequences were obtained from UniProt and their sequences aligned using Clustal Omega^63^. AlphaFold2 multimer^64^ was used through the implementation in ChimeraX^60^. Six copies of the selected CBS Bateman domains were used as the input and the top scored model was used for structural analysis. All lower scored models were essentially identical to the top scorer for all orthologues.

### Data availability

Structures and EM maps of CBS basal state (EMDB-XXXX, PDB-XXXX, EMDB-XXXXX, PDB-XXXX, EMDB-XXXX, PDB-XXXX, EMDB-XXXXX, PDB-XXXX), degraded CBS tetramer (EMDB-XXXXX, PDB-XXXX), CBS+SAM global activated state (EMDB-XXXXX, PDB-XXXX), as well as EM maps for CBS+SAM regulatory domains activated state (EMDB-XXXXX, EMDB-XXXX) and CBS+SAM catalytic domain activated state (EMDB-XXXXX), have been deposited to the EMDB and PDB databases.

## Abbreviations

CBS: cystathionine beta-synthase
PLP: pyridoxal 5’ phosphate
SAM: s-adenosyl-L-methionine
SAH: s-adenosyl-L-homocysteine
ITC: isothermal titration calorimetry
CD: catalytic domain
RD: regulatory domain
HCU: homocystinuria

## Acknowledgements

We thank all members of the SGC Oxford Biotechnology team for their molecular biology support. We thank the Oxford Particle Imaging Centre (OPIC) electron microscopy facility for grid screening and data collection. We acknowledge the Diamond Light Source for access and support to the UK’s Electron Bio-imaging Centre (eBIC, under BAG proposal EM20223) funded by the Wellcome Trust, MRC, and BBRSC. We specifically want to thank James Gilchrist and Yuriy Chaban for their assistance. We also want to thank Laura Diez-Siez for her help in the initial EM screening and data collection. Additionally, we thank Johan Turkenburg, Sam Hart, and Jamie Blaza for their assistance in collecting Glacios data at York Structural Biology Laboratory (YSBL). The YSBL is funded by the BBSRC, the Wellcome Trust (grant number 206161/Z/17/Z), Tony Wild, and the Wolfson Foundation. We also wish to thank Brian Marsden and Chris Sluman for their bioinformatics support. The Structural Genomics Consortium is a registered charity (Number 1097737) that receives funds from AbbVie, Bayer Pharma AG, Boehringer Ingelheim, Canada Foundation for Innovation, Eshelman Institute for Innovation, Genome Canada, Innovative Medicines Initiative (EU/EFPIA) [ULTRA-DD grant no. 115766], Janssen, Merck & Co., Novartis Pharma AG, Ontario Ministry of Economic Development and Innovation, Pfizer, São Paulo Research Foundation-FAPESP, Takeda, and Wellcome Trust [092809/ Z/10/Z]. Initial work on this project was funded by a Wellcome Trust Pathfinder Award to W.W.Y. T.J.M. received cryo-EM training through Wellcome/MRC funded program (218785/Z/19/Z), was a recipient of a Pump-Priming Award from the Medical Sciences Division, University of Oxford, and is funded by a HCU Network North America and Australia Research grant.

## Author contributions

T.J.M. and W.W.Y. designed the experiments. T.J.M. expressed and purified all CBS constructs, carried out biochemical experiments, screened, and collected EM data, analyzed, and refined all CBS structures. H.J.B. did initial screening and collected EM data. C.S.D. carried out initial CBS construct cloning, expression testing, and optimization. A.B. assisted in EM processing of the CBS^FL^ basal state. T.J.M. and W.W.Y. carried out the data analysis and wrote the manuscript with contributions from all authors.

---

**Extended Data Fig. 1.**
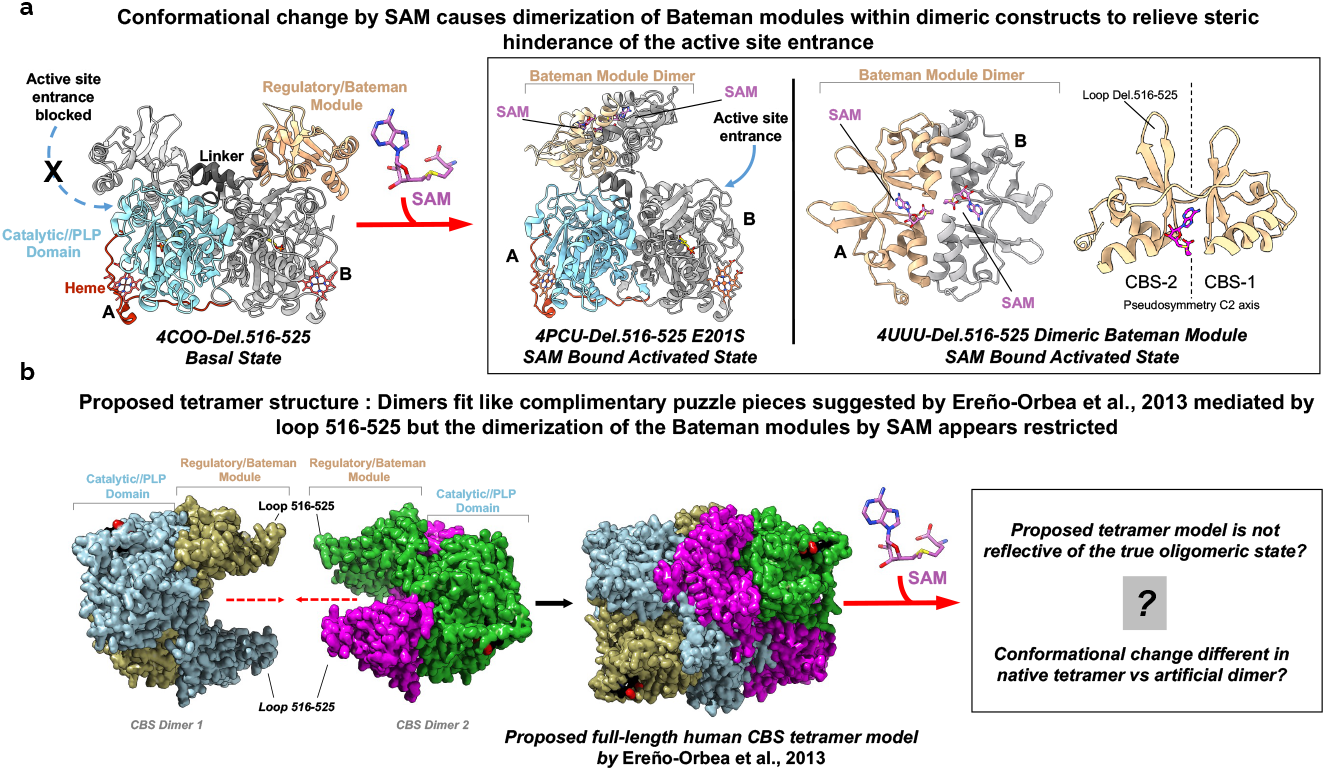
Previous crystal structures of CBS^Δ516–525^ without and with SAM, and the proposed tetrameric state. **a**, Crystal structures of dimeric CBS^Δ516–525^ without and with SAM demonstrating a domain un-swapping and dimerization of the Bateman modules in the SAM bound activated state. In the basal state CBS is a domain swapped dimer where the neighboring subunits regulatory/Bateman module sits atop the catalytic site entrance. Two crystal structures, one of a mutant and another of the loop-deleted regulatory domain alone, show that SAM binding causes disassociation of the regulatory domain from the catalytic domain resulting in dimerization to form a Bateman module dimer. **b**, Ereño-Orbea et al., 2013 suggested that CBS oligomerizes by clasping interactions of loop 516-525 from two domain swapped dimers to form an intimate tetrameric particle. However, this proposed architecture is not compatible with the known crystal structures of the SAM bound activated state. This suggests that the proposed tetrameric state is incorrect, or the known conformational change is different within the context of the full-length CBS tetramer.

**Extended Data Fig. 2.**
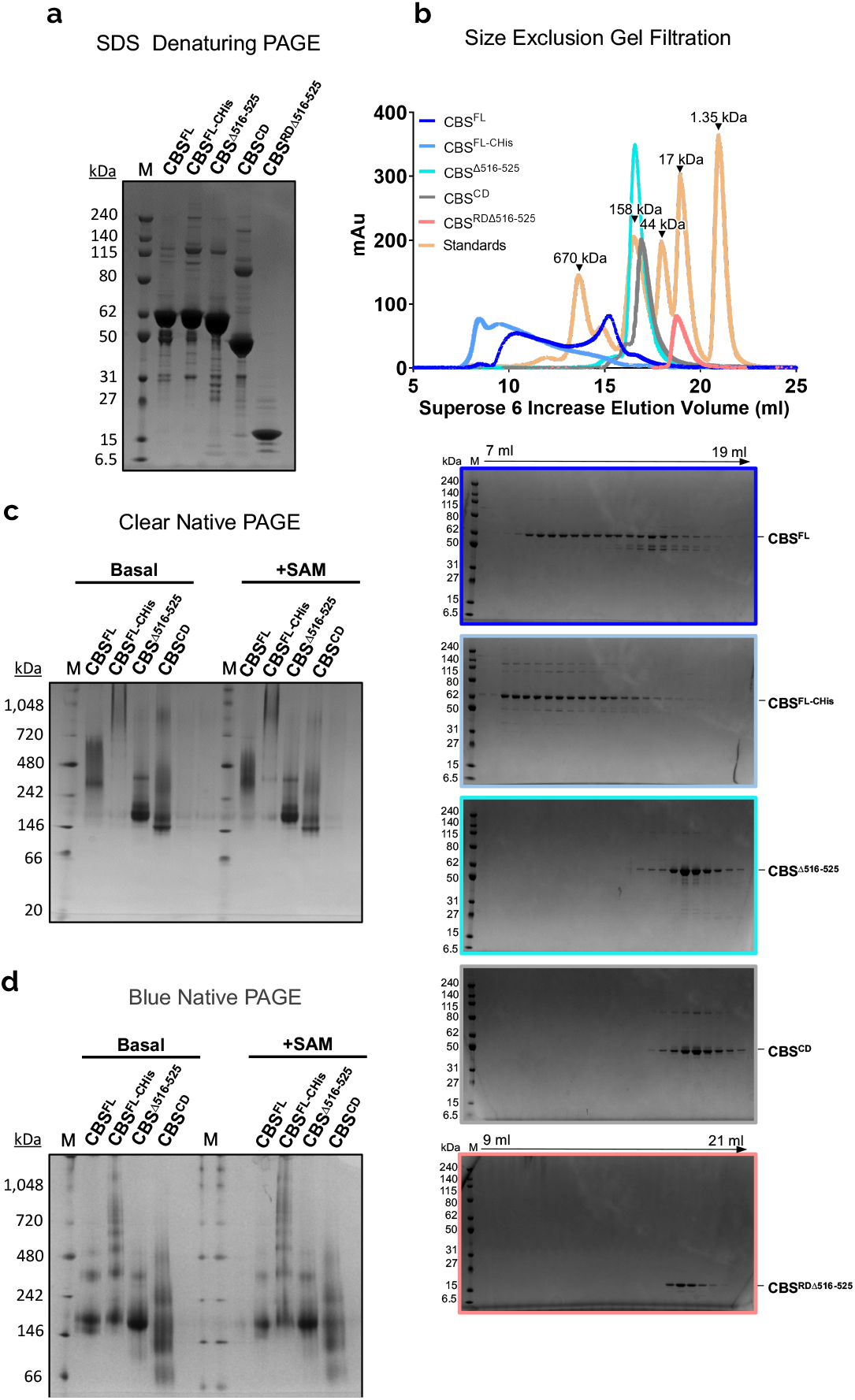
Analysis of the oligomeric states of different CBS constructs. **a**, Coomassie stained SDS-PAGE of the five CBS constructs used in this study. **b**, Superose 6 gel filtration chromatograph and Coomassie stained SDS-PAGE gels of each CBS constructs. n = 1. **c**, Representative clear-native PAGE of each CBS construct without and with 1 mM SAM. *n* = 2 technical repeats. **d,** Representative blue-native PAGE of each CBS construct without and with 1 mM SAM. *n* = 2 technical repeats.

**Extended Data Fig. 3.**
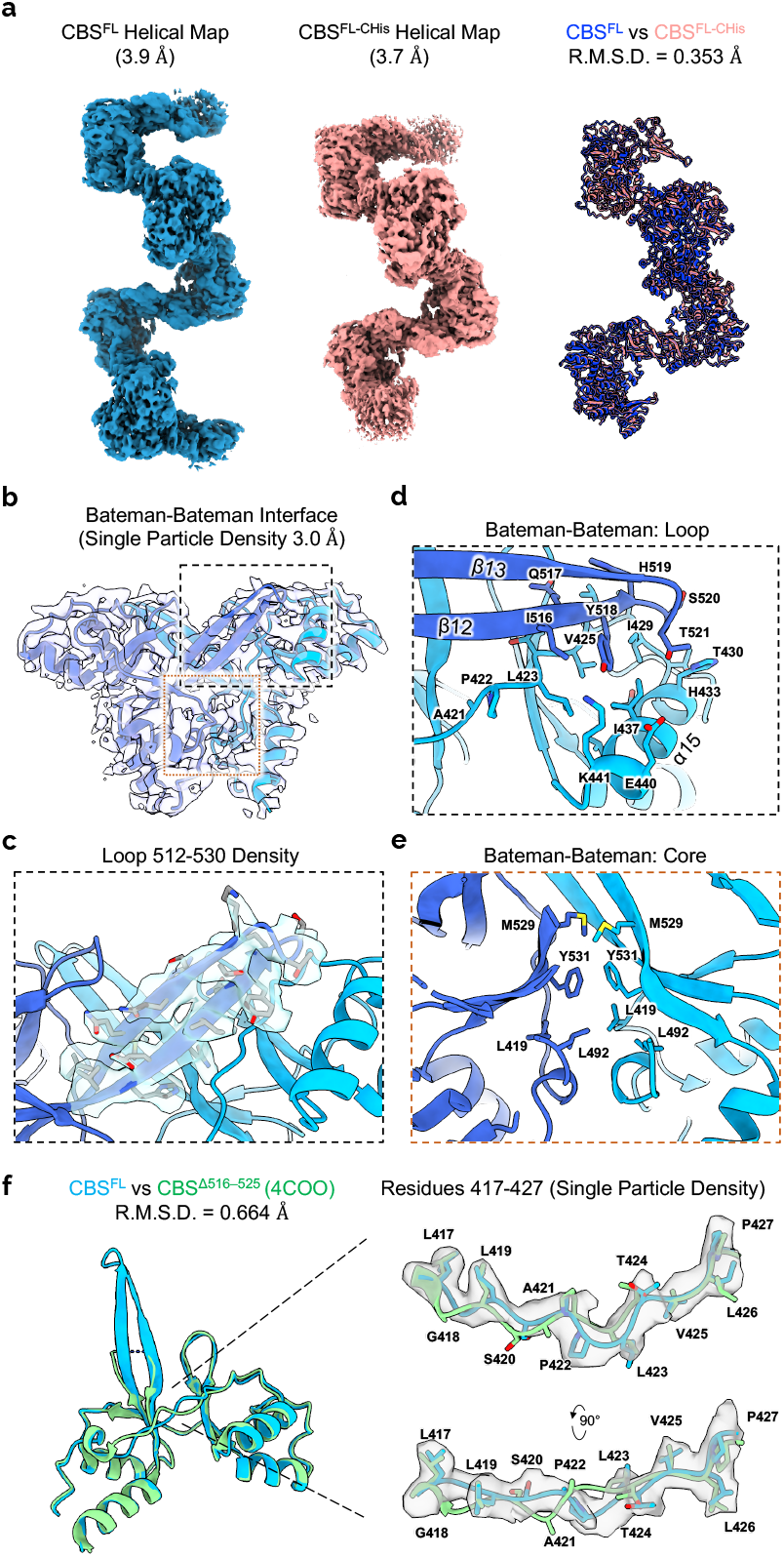
Key residues are involved in the polymerization of human CBS. **a**, Cryo-EM maps of both CBS^FL^ and CBS^FL-CHis^ determined using helical reconstruction. Alignment of the resulting refined atomic models show very little difference. **b**, Density and model fit at the Bateman-Bateman interface of the 3.0 Å resolution single particle map of CBS^FL-CHis^. **c**, Close up of the cryo-EM density and residues of the loop 512-530 at the Bateman-Bateman interface. **d**, Close up of residues involved in interactions at the loop 516-525 of the Bateman-Bateman interface. **e**, Close up of residues involved in hydrophobic interactions at the core of the Bateman-Bateman interface. **f**, Structural alignment of the regulatory domains from the cryo-EM map of filamentous CBS^FL-CHis^ and crystal structure of dimeric CBS^Δ516–525^ (4COO). Interactions due to filamentation result in a shift of the residues 417-423.

**Extended Data Fig. 4.**
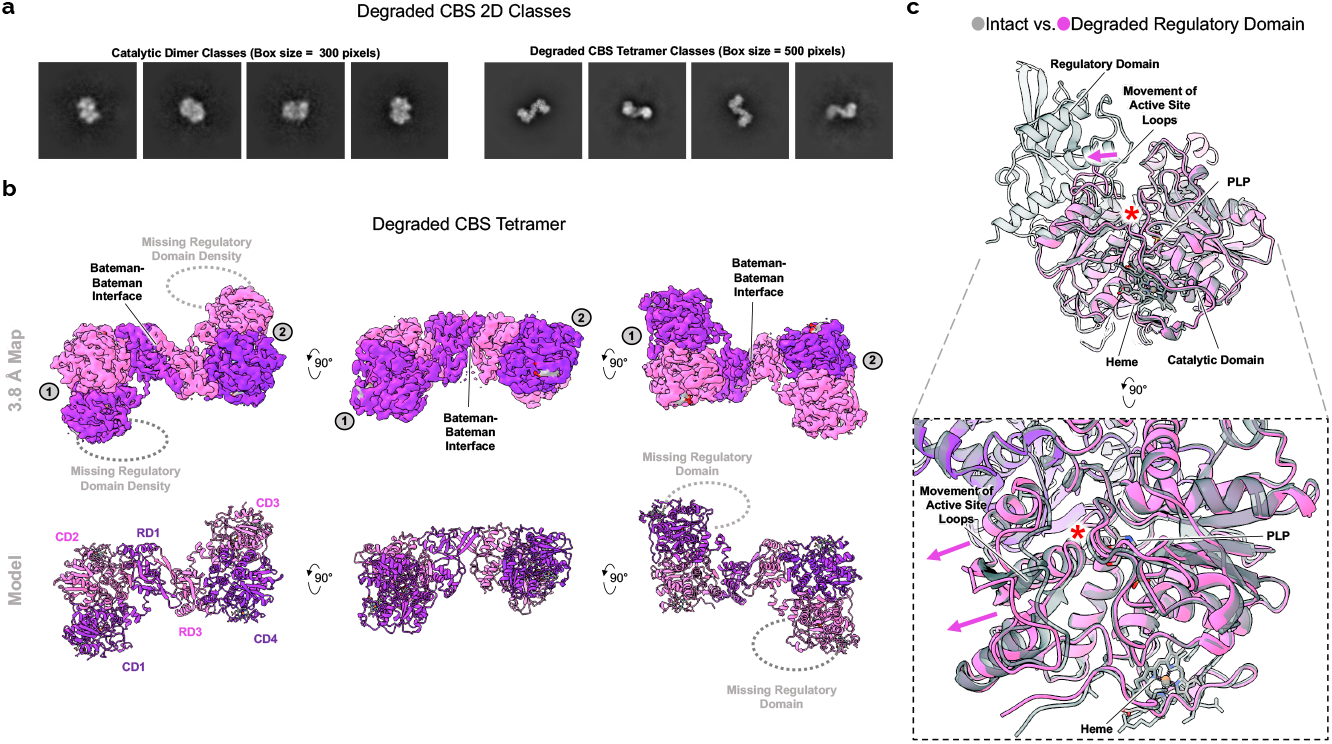
CBS^FL^ degrades into a tetramer and dimer. **a**, Example 2D classes representative of degraded CBS catalytic domain dimer and degraded tetrameric CBS. **b**, Multiple views of the 3.8 Å resolution map and model of tetrameric degraded CBS. The tetramer is formed by Bateman-Bateman interactions of two CBS heterodimers consisting of one full-length protomer (residues 42-548) and one degraded protomer without the regulatory domain (residues 42-398). The regulatory domain has been proteolyzed at the flexible linker (residues 382-411) between it and the catalytic domain. **c**, Structural alignment of intact CBS^FL^ vs degraded CBS. The absence of the regulatory domain allows the active site loops to open and increases accessibility to the active site denoted as the red asterisk.

**Extended Data Fig. 5.**
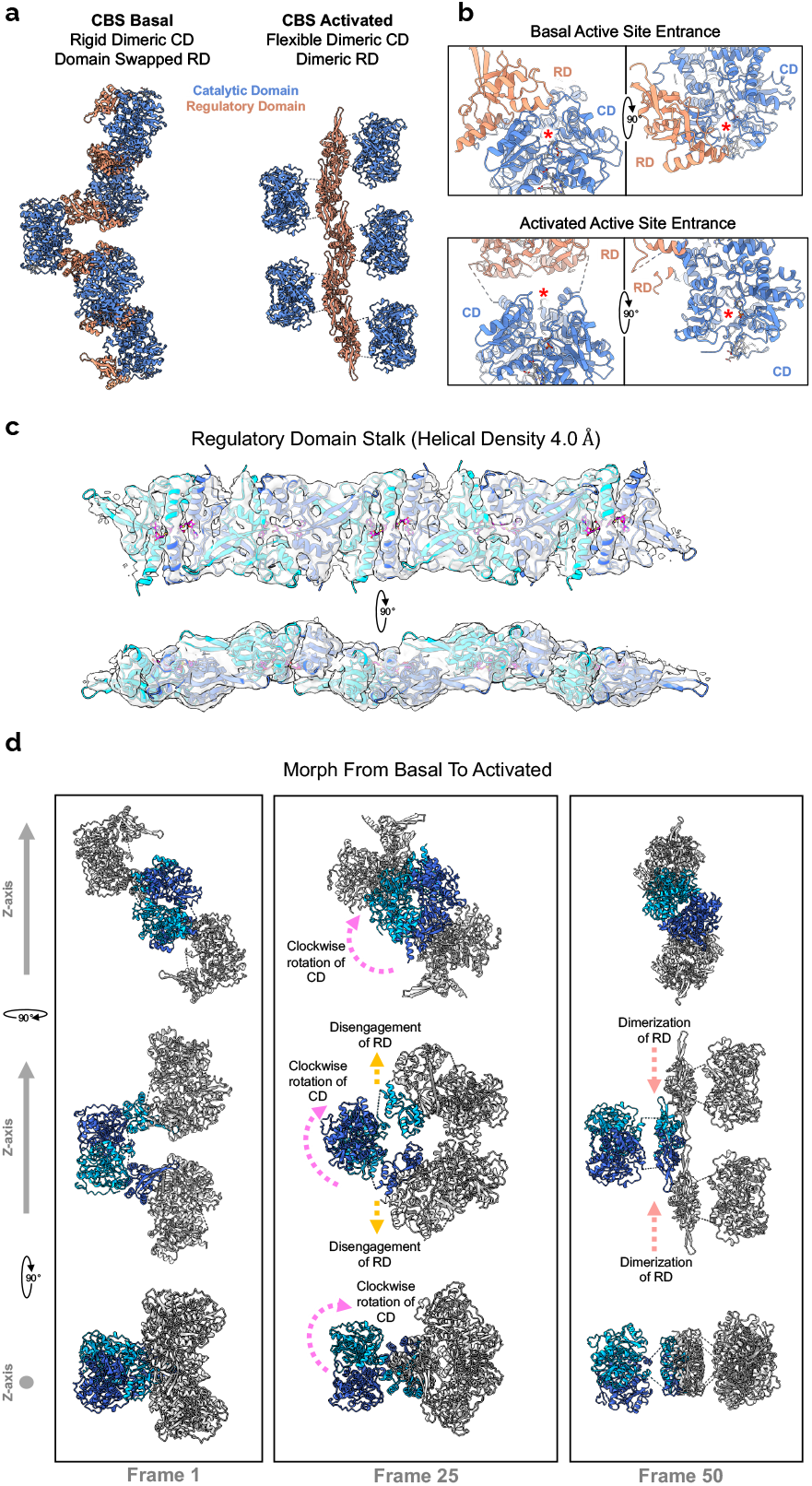
The allosteric change initiated by SAM binding is compatible with the filamentous architecture of human CBS. **a**, Structures of the basal and SAM bound activated state showing the relative positions of the catalytic and regulatory domains. **b**, Closeup of the active site entrance in both basal and activated states. The active site entrance is denoted as a red asterisk. **c**, Cryo-EM density, and model fit of the central regulatory domain stalk of the activated state. **d**, A morph of one turn of the CBS filament from the basal to activated state. Both models when aligned to the central Z-axis suggest that the conformational change due to SAM requires rotation of the catalytic domain along with disengagement and dimerization of the regulatory domains.

**Extended Data Fig. 6.**
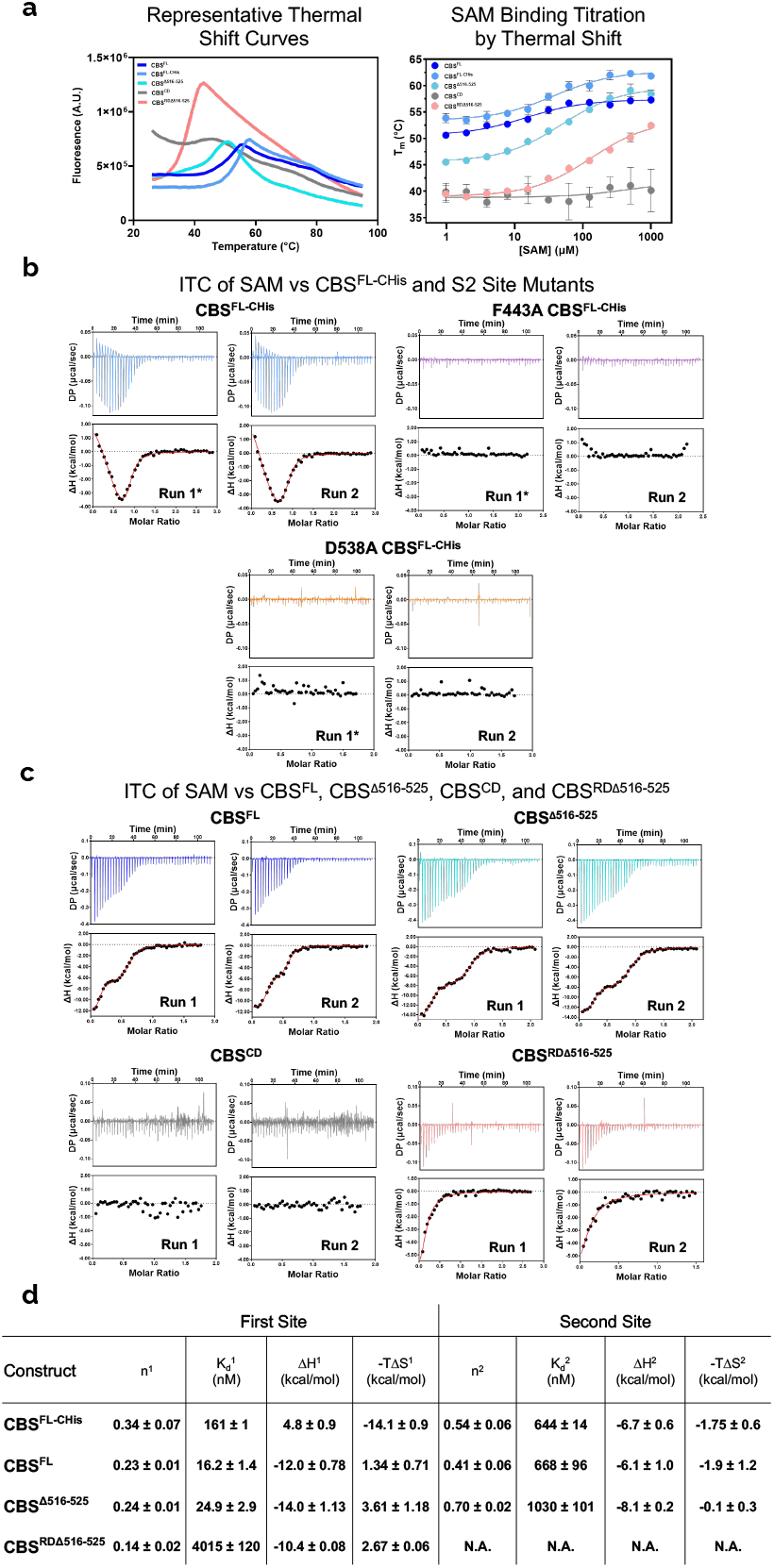
ITC analysis of SAM titrations into constructs representative of the domain arrangement of CBS. **a**, Representative thermal shift curves and SAM titrations for CBS^FL-CHis^, CBS^FL^, CBS^Δ516–525^, CBS^CD^, and CBS^RDΔ516–525^ **b**, Replicate ITC titrations of SAM against wild-type, F443A, and D538A CBS^FL-CHis^. Run 1 is presented in Figure 3 and is shown here for comparison purposes only. **c**, Replicate ITC titrations of SAM against CBS^FL^, CBS^Δ516–525^, CBS^CD^, and CBS^RDΔ516–525^. **d**, Calculated ITC parameters for SAM binding against CBS^FL-CHis^, CBS^FL^, CBS^Δ516–525^, and CBS^RDΔ516–525^. Binding parameters were determined by fitting to one or two site binding appropriately. Values are means and ± s.d. of *n* = 2 technical repeats.

**Extended Data Fig. 7.**
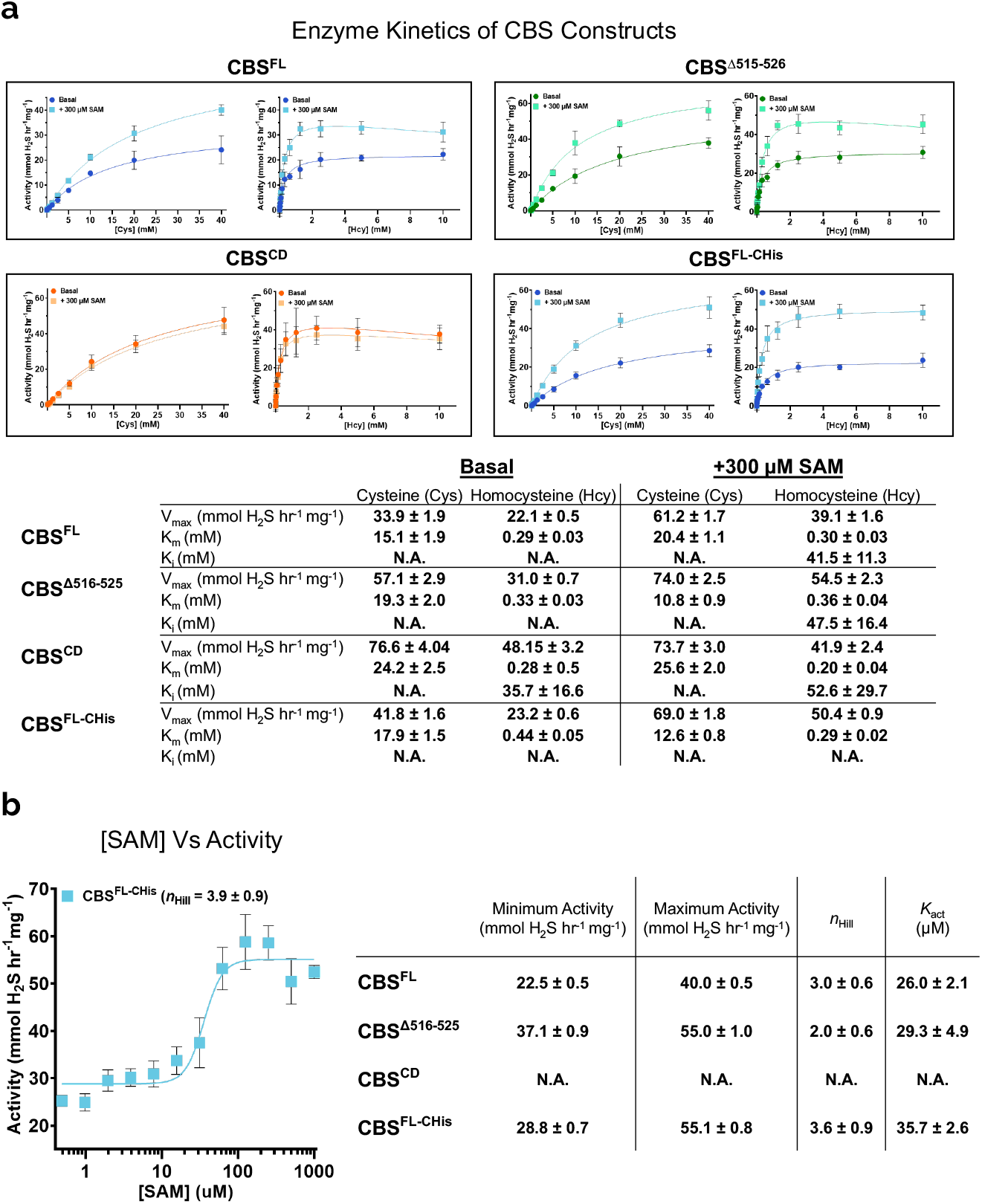
The enzyme activity and SAM activation of the CBS constructs in this study. **a**, Enzyme kinetics of CBS^FL^, CBS^Δ516–525^, CBS^CD^, and CBS^FL-CHis^ without (Basal) and with SAM (+ 300 μM SAM). Mean values and error bars are ± s.d. of *n* = at least 6 technical repeats. **b**, H2S producing activity of CBS^FL-CHis^ in response to increasing amounts of SAM. Values are means and error bars are ± s.d. of *n* = at least 4 technical repeats **c**, Fitted values for the allosteric activation by SAM of CBS^FL^, CBS^Δ516–525^, CBS^CD^, and CBS^FL-CHis^. Values are means and ± s.d. of *n*= at least 4 technical repeats

**Extended Data Fig. 8.**
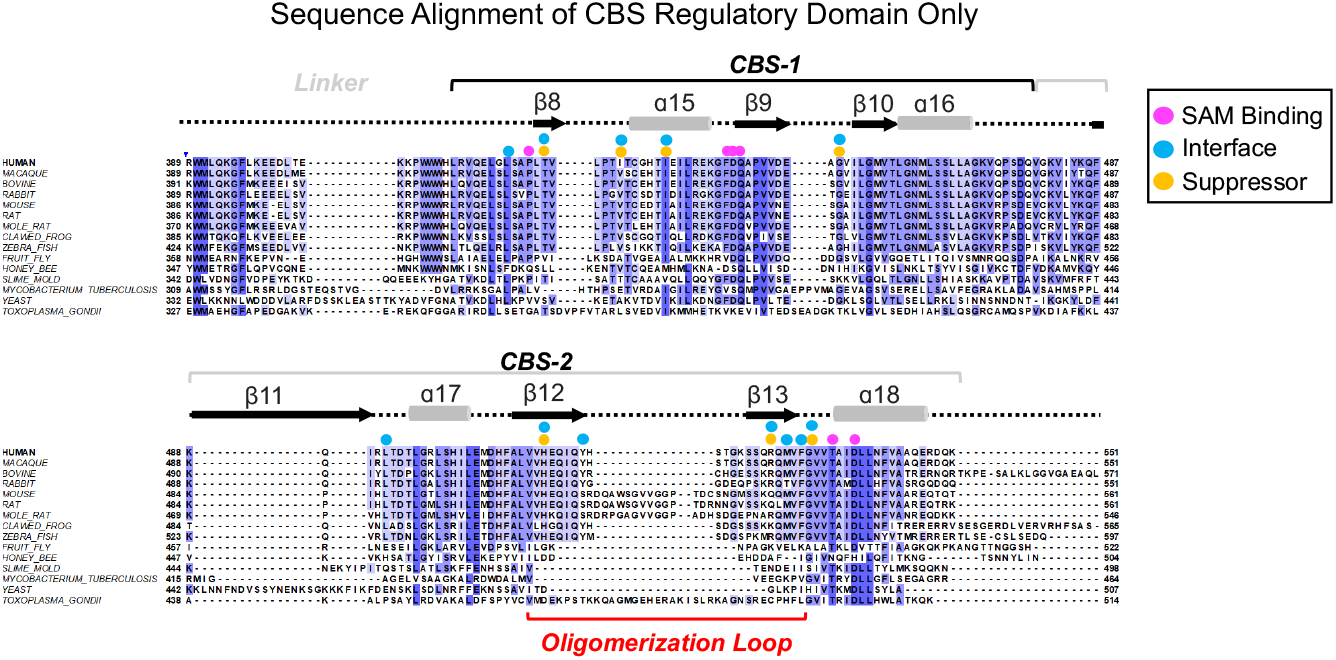
Sequence alignment of the regulatory domain of CBS enzymes. Sequence alignment of the regulatory domain only of various CBS enzymes. The *CBS-1* and *CBS-2* motifs are indicated.

**Extended Data Fig. 9.**
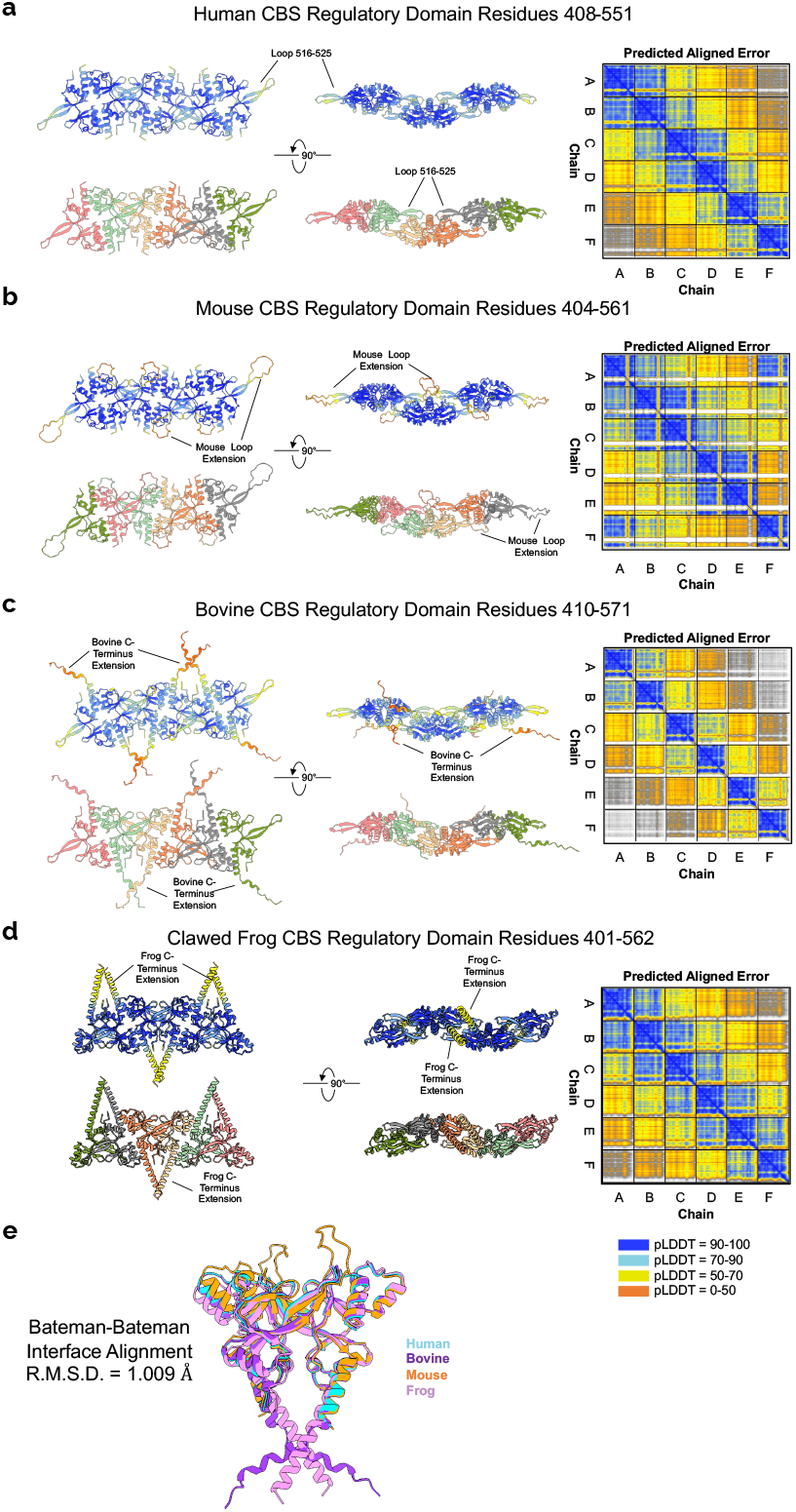
AlphaFold predicts SAM responsive CBS enzymes polymerise as filaments. **a**, AlphaFold prediction using six copies of the human CBS regulatory domain (residues 408-551) shows a filament architecture identical to the cryo-EM model of the activated state. **b**, AlphaFold prediction using six copies of the mouse CBS regulatory domain (residues 404-561) shows a filament architecture similar to the cryo-EM model of the human CBS activated state. **c**, AlphaFold prediction using six copies of the bovine CBS regulatory domain (residues 410-571) shows a filament architecture similar to the cryo-EM model of the human CBS activated state. **d**, AlphaFold prediction using six copies of the African clawed frog CBS regulatory domain (residues 401-562) shows a filament architecture similar to the cryo-EM model of the human CBS activated state. **e**, Structural alignment of the Bateman-Bateman interface from the AlphaFold predicted structures of human, mouse, bovine and frog CBS.

**Extended Data Fig. 10.**
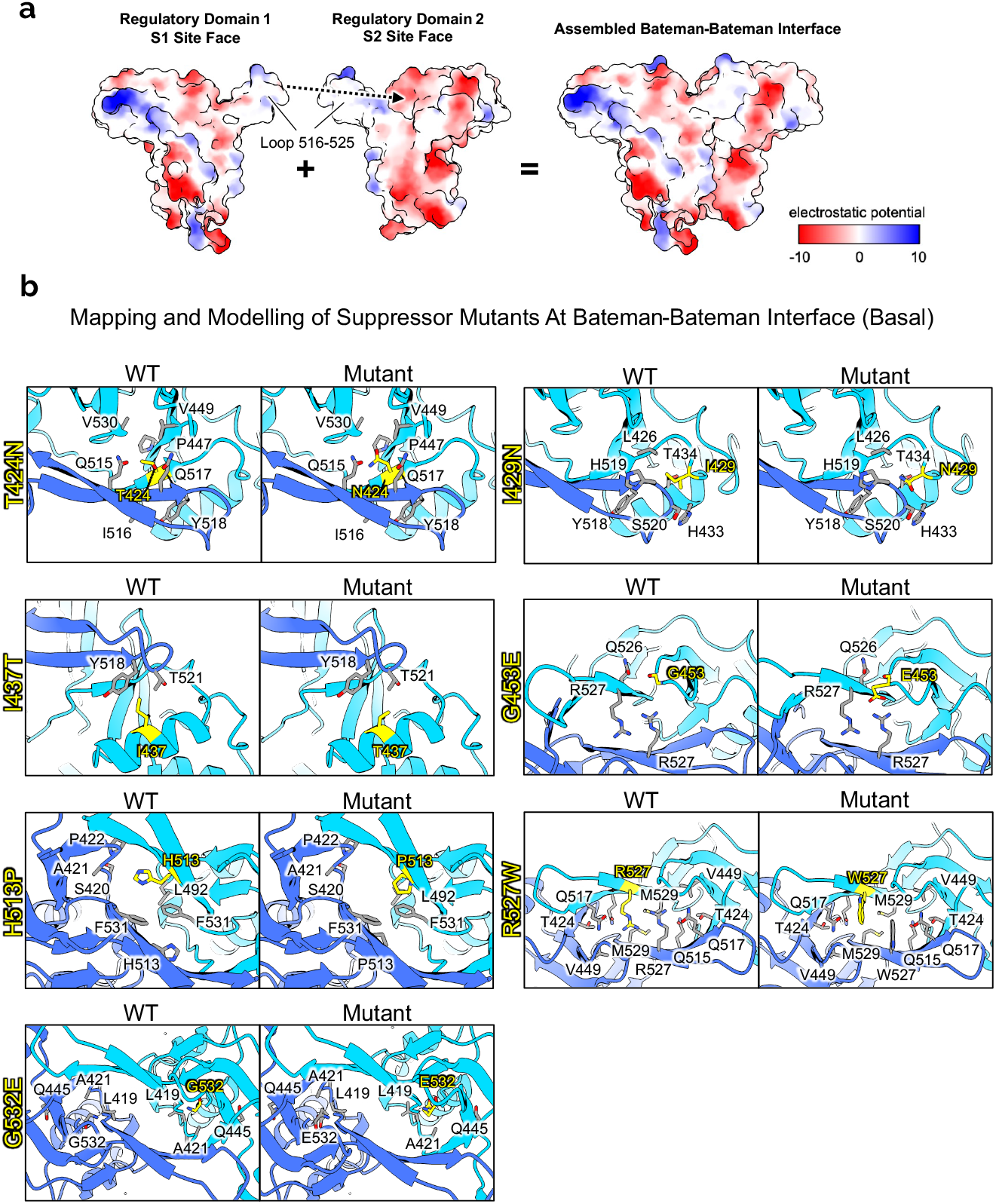
Hydrophobic interaction of CBS regulatory domains and analysis of suppressor mutations. **a**, Surface electrostatic potential of the CBS regulatory domain shows that assembly is mainly driven by hydrophobic interactions mediated by the loop 516-525. The interface involves the S2 site face of both regulatory domains **b**, Structural analysis of the seven reported HCU suppressor mutations mapped and statically mutated. All are located at the Bateman-Bateman interface and are predicted to alter oligomerization.

**Extended Data table 1.** Cryo-EM data collection, refinement, and validation statistics.

|  | Helical basal<br>state with His<br>tag<br>(EMDB-XXXXX)<br>(PDB-XXXX) | SPA basal<br>state with His<br>tag<br>(EMDB-XXXXX)<br>(PDB XXXX) | Helical basal state<br>(EMDB-XXXXX)<br>(PDB XXXX) | SPA basal state<br>(EMDB-XXXXX)<br>(PDB XXXX) | Tetramer basal<br>state<br>(EMDB-XXXXX)<br>(PDB XXXX) | Global helical<br>activated state<br>(EMDB-XXXXX)<br>(PDB XXXX) | Focused helical<br>activated state<br>(EMDB-XXXXX) | Focused SPA<br>activated state<br>regulatory domain<br>(EMDB-XXXXX) | Focused SPA<br>activated state<br>catalytic domain<br>(EMDB-XXXXX) |
| --- | --- | --- | --- | --- | --- | --- | --- | --- | --- |
| **Data collection and processing** |  |  |  |  |  |  |  |  |  |
| Magnification | 75,000 |  | 150,000 |  |  | 81,000 |  |  |  |
| Voltage (kV) | 300 |  | 200 |  |  | 300 |  |  |  |
| Electron exposure (e<sup>-</sup>/Å<sup>2</sup>) | 37.85 |  | 50.00 |  |  | 39.96 |  |  |  |
| Defocus range (µm) | -0.9 to -3.0 |  | -1.4 to -2.0 |  |  | -0.9 to -3.0 |  |  |  |
| Pixel size (Å) | 1.083 |  | 0.934 |  |  | 1.06 |  |  |  |
| Symmetry imposed | D1 | C2 | D1 | C2 | C2 | D1 | D1 | D1 | D1 |
| Initial particle images (no.) | 239,739 | 760,869 | 749,213 | 1,190,611 | 1,190,611 | 5,240,414 | 5,240,414 | 5,240,414 | 5,240,414 |
| Final particle images (no.) | 76,663 | 113,271 | 89,761 | 96,182 | 67,901 | 425,261 | 425,261 | 425,261 | 425,261 |
| Map resolution (Å) | 3.7 | 3.0 | 3.9 | 3.8 | 3.8 | 4.0 | 4.1 | 4.1 | 8.3 |
| FSC threshold | 0.143 | 0.143 | 0.143 | 0.143 | 0.143 | 0.143 | 0.143 | 0.143 | 0.143 |
| Map resolution range (Å) | 3.0-6.0 | 2.6-6.0 | 3.3-7.3 | 3.2-7.1 | 3.4-7.7 | 3.3-11.0 | 3.5-13.0 | 4.0-7.0 | 6.7-10.3 |
| **Refinement** |  |  |  |  |  |  |  |  |  |
| Initial model used (PDB code) | 4COO | 4COO | 4COO | 4COO | 4COO | 4UUU, 4PCU |  |  |  |
| Model resolution (Å) | 3.9 | 3.3 | 4.3 | 3.9 | 4.0 | 13.5 |  |  |  |
| FSC threshold | 0.5 | 0.5 | 0.5 | 0.5 | 0.5 | 0.5 |  |  |  |
| Model resolution range (Å) | n/a | n/a | n/a | n/a | n/a | n/a |  |  |  |
| Map sharpening *B* factor (Å<sup>2</sup>) | -90 | -80 | - | -90 | -80 | -70 | -80 | -110 | -90 |
| Model composition |  |  |  |  |  |  |  |  |  |
| Nonhydrogen atoms | 31939 | 23928 | 39917 | 15956 | 13590 | 50900 |  |  |  |
| Protein residues | 4065 | 3042 | 5079 | 2028 | 1728 | 4970 |  |  |  |
| Ligands | 8 | 6 | 10 | 4 | 6 | 20 |  |  |  |
| *B* factors (Å<sup>2</sup>) |  |  |  |  |  |  |  |  |  |
| Protein | 68.04 | 40.53 | 25.87 | 81.32 | 92.56 | 178.67 |  |  |  |
| Ligand | 98.88 | 51.94 | 81.41 | 87.17 | 88.80 | 168.47 |  |  |  |
| R.m.s. deviations |  |  |  |  |  |  |  |  |  |
| Bond lengths (Å) | 0.004 | 0.003 | 0.003 | 0.003 | 0.005 | 0.002 |  |  |  |
| Bond angles (°) | 0.701 | 0.602 | 0.645 | 0.631 | 0.762 | 0.566 |  |  |  |
| Validation |  |  |  |  |  |  |  |  |  |
| MolProbity score | 1.95 | 1.66 | 1.90 | 1.82 | 1.96 | 2.35 |  |  |  |
| Clashscore | 14.28 | 10.02 | 11.65 | 10.05 | 14.00 | 33.82 |  |  |  |
| Poor rotamers (%) | 0.38 | 0.23 | 0.35 | 0.53 | 0.14 | 0.00 |  |  |  |
| Ramachandran plot |  |  |  |  |  |  |  |  |  |
| Favored (%) | 95.77 | 97.28 | 95.38 | 95.67 | 95.57 | 95.10 |  |  |  |
| Allowed (%) | 4.23 | 2.59 | 4.62 | 4.33 | 4.43 | 4.90 |  |  |  |
| Disallowed (%) | 0.00 | 0.00 | 0.00 | 0.00 | 0.00 | 0.20 |  |  |  |

## Notes

### Competing Interest Statement

The authors have declared no competing interest.

