## Supplementary Information for "Architecture and regulation of filamentous human cystathionine beta-synthase"

**Table of Contents**

**Supplementary Fig. 1:** Helical cryo-EM data processing of basal state CBS$^{FL-CHis}$…………..2
**Supplementary Fig. 2:** Single particle cryo-EM data processing of basal state CBS$^{FL-CHis.}$…3
**Supplementary Fig. 3:** Helical cryo-EM data processing of basal state CBS$^{FL}$……..………..4
**Supplementary Fig. 4:** Single particle cryo-EM data processing of basal state CBS$^{FL}$ …….5
**Supplementary Fig. 5:** Helical cryo-EM data processing of activated state CBS$^{FL-CHis}$ ..…...6
**Supplementary Fig. 6:** Further cryo-EM data processing of activated state CBS$^{FL-CHis}$ …….7
**Supplementary Fig. 7:** Source images of Extended Data Fig. 2a-d……………………..…….8
**Supplementary Movie 1:** Morph of the Bateman-Bateman interface from the basal and SAM bound activated structure of CBS ……………………………………………………..……9
**Supplementary Movie 2:** Morph of one full turn of the basal and SAM bound activated structures of CBS…...………………………………………………………………………..……9

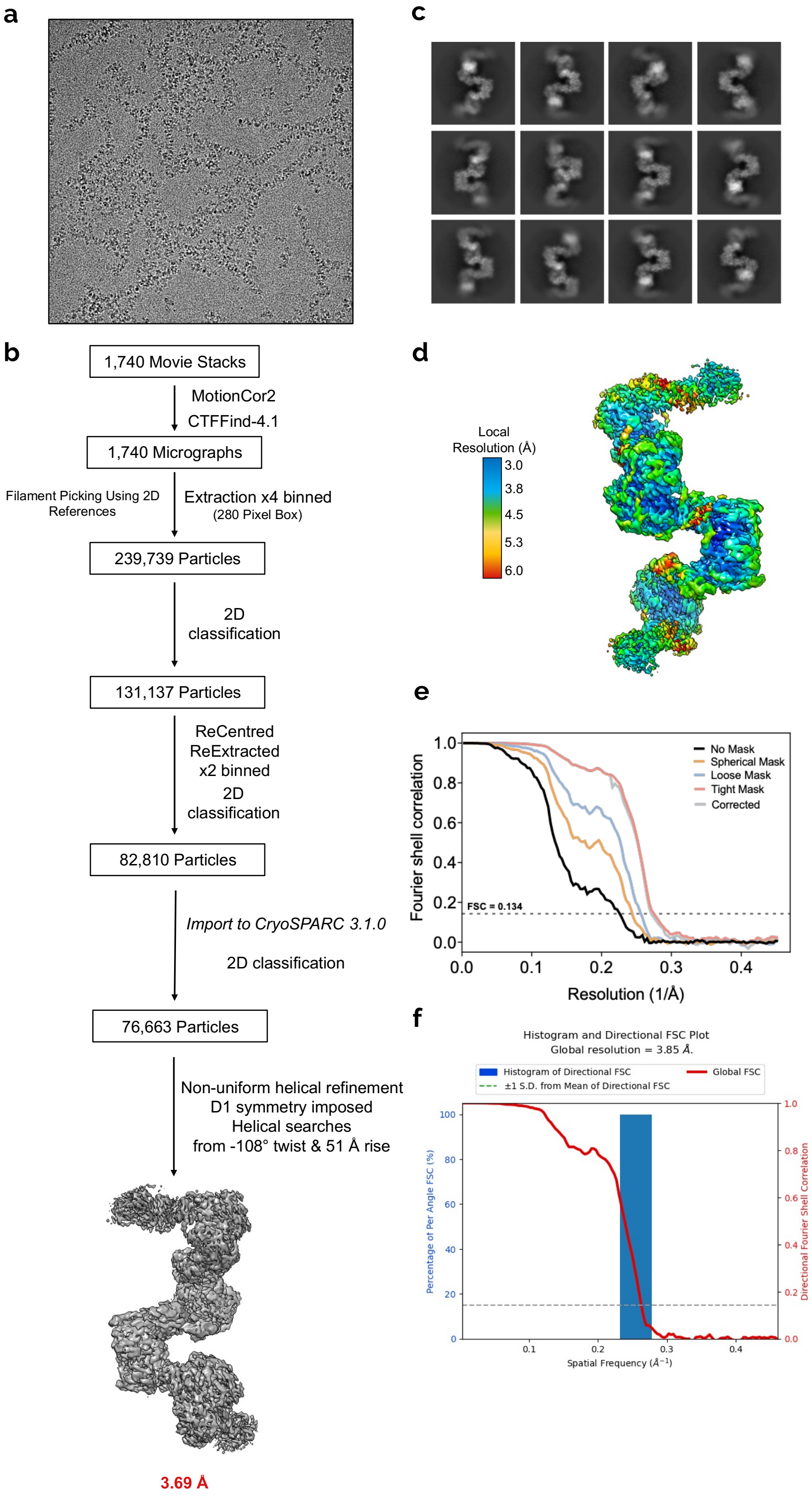


**Supplementary Fig. 1 | Helical cryo-EM data processing of basal state CBS$^{FL\text{-}CHis}$. a**, Representative Falcon 3 micrograph of CBS$^{FL\text{-}CHis}$. **b**, Processing flow chart of CBS$^{FL\text{-}CHis}$ in the basal state. **c**, Representative helical 2D classes of CBS$^{FL\text{-}CHis}$. **d**, Local resolution variation of the 3.7 Å helical CBS$^{FL\text{-}CHis}$ basal state map. **e**, Fourier shell correlation (FSC) curve. **f**, Directional FSC plot of the helical CBS$^{FL\text{-}CHis}$ map.

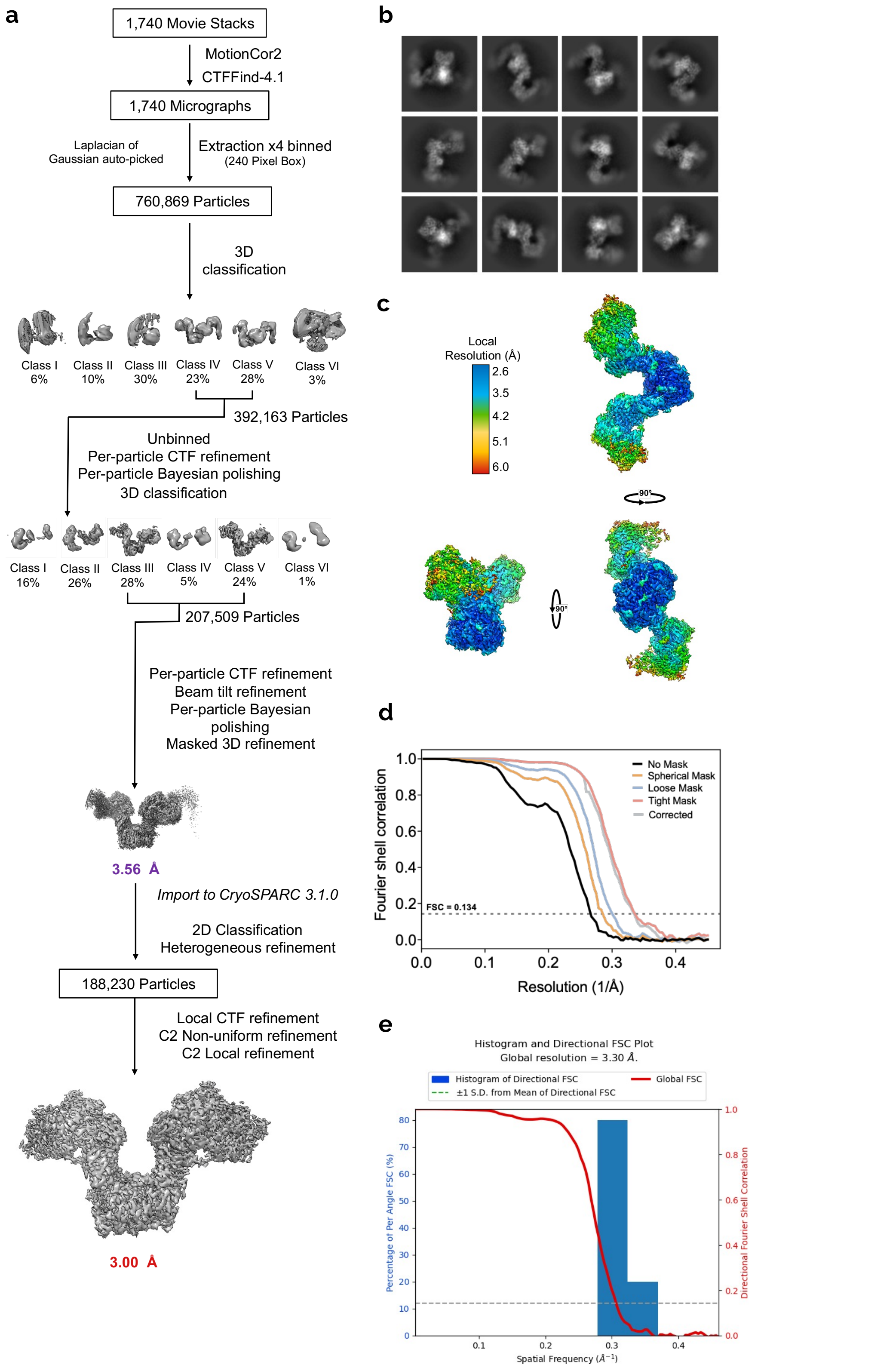


**Supplementary Fig. 2 | Single particle cryo-EM data processing of basal state CBS$^{FL\text{-}CHis}$.** **a**, Processing flow chart of CBS$^{FL\text{-}CHis}$ in the basal state. **b**, Representative 2D classes of CBS$^{FL\text{-}CHis}$. **c**, Local resolution variation of the 3.0 Å CBS$^{FL\text{-}CHis}$ basal state map. **d**, Fourier shell correlation (FSC) curve. **e**, Directional FSC plot of the 3.0 Å CBS$^{FL\text{-}CHis}$ map.

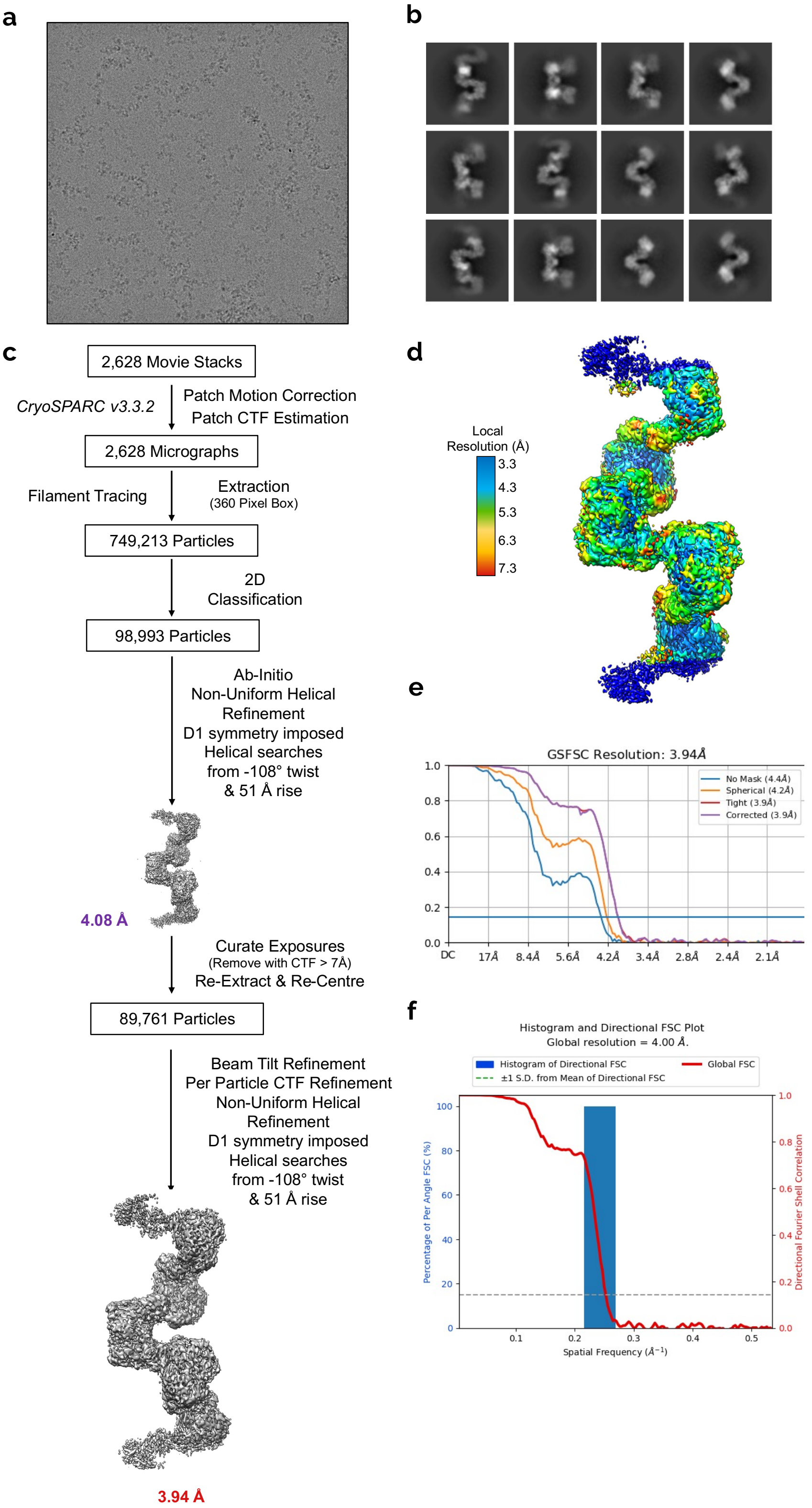


**Supplementary Fig. 3 | Helical cryo-EM data processing of basal state CBS$^{FL}$. a**, Representative Falcon 4 micrograph of CBS$^{FL}$. **b**, Processing flow chart of CBS$^{FL}$ in the basal state. **c**, Representative helical 2D classes of CBS$^{FL}$. **d**, Local resolution variation of the 3.9 Å helical CBS$^{FL}$ basal state map. **e**, Fourier shell correlation (FSC) curve. **f**, Directional FSC plot of the 3.9 Å helical CBS$^{FL}$ map

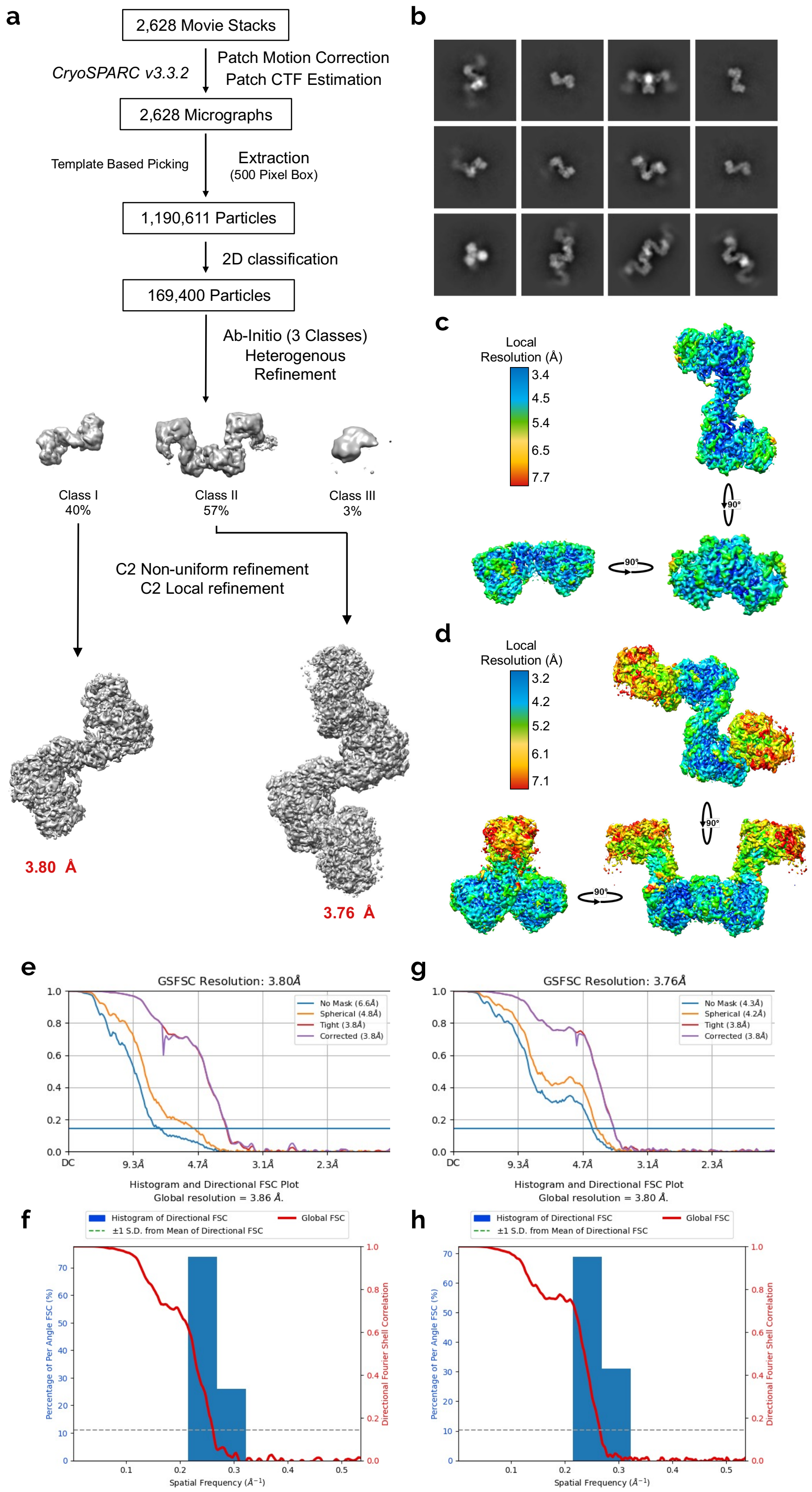


**Supplementary Fig. 4 | Single particle cryo-EM data processing of basal state CBS$^{FL}$. a**, Processing flow chart of CBS$^{FL}$ in the basal state. **b**, Representative 2D classes of CBS$^{FL}$. **c, d,** Local resolution variation of the two CBS$^{FL}$ basal state maps. **e, f,** Fourier shell correlation (FSC) curves. **f, h,** Directional FSC plots of the two CBS$^{FL}$ maps.

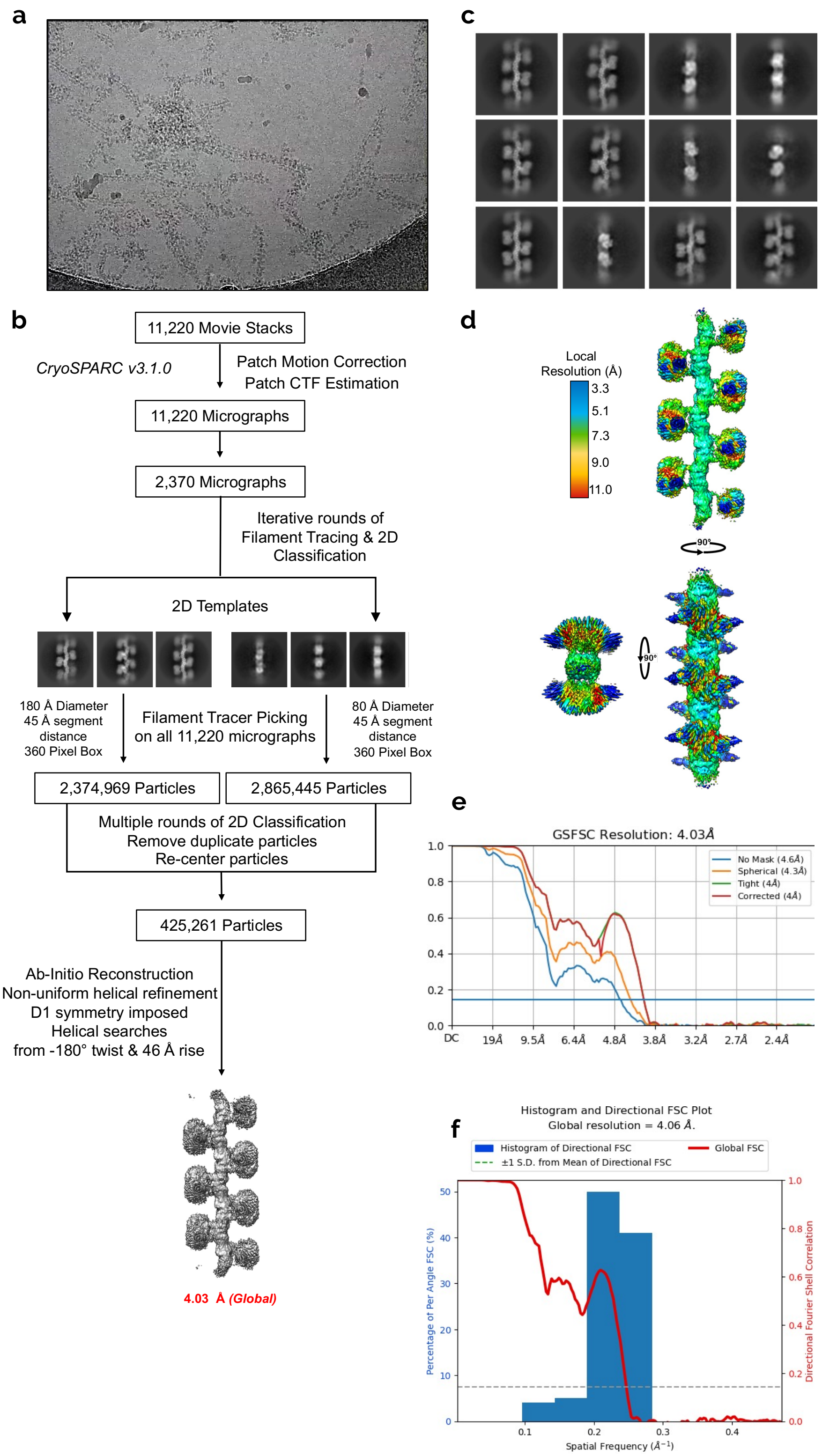


**Supplementary Fig. 5 | Helical cryo-EM data processing of activated state CBS$^{FL\text{-}CHis}$. a**, Representative K3 micrograph of CBS$^{FL\text{-}CHis}$ in the presence of SAM. **b**, Processing flow chart of CBS$^{FL\text{-}CHis}$ in the activated, SAM bound, state. **c**, Representative helical 2D classes of CBS$^{FL\text{-}CHis}$ bound to SAM. **d**, Local resolution variation of the 4.0 Å helical CBS$^{FL\text{-}CHis}$ activated state map. **e**, Fourier shell correlation (FSC) curve. **f**, Directional FSC plot of the helical CBS$^{FL\text{-}CHis}$ plus SAM map.

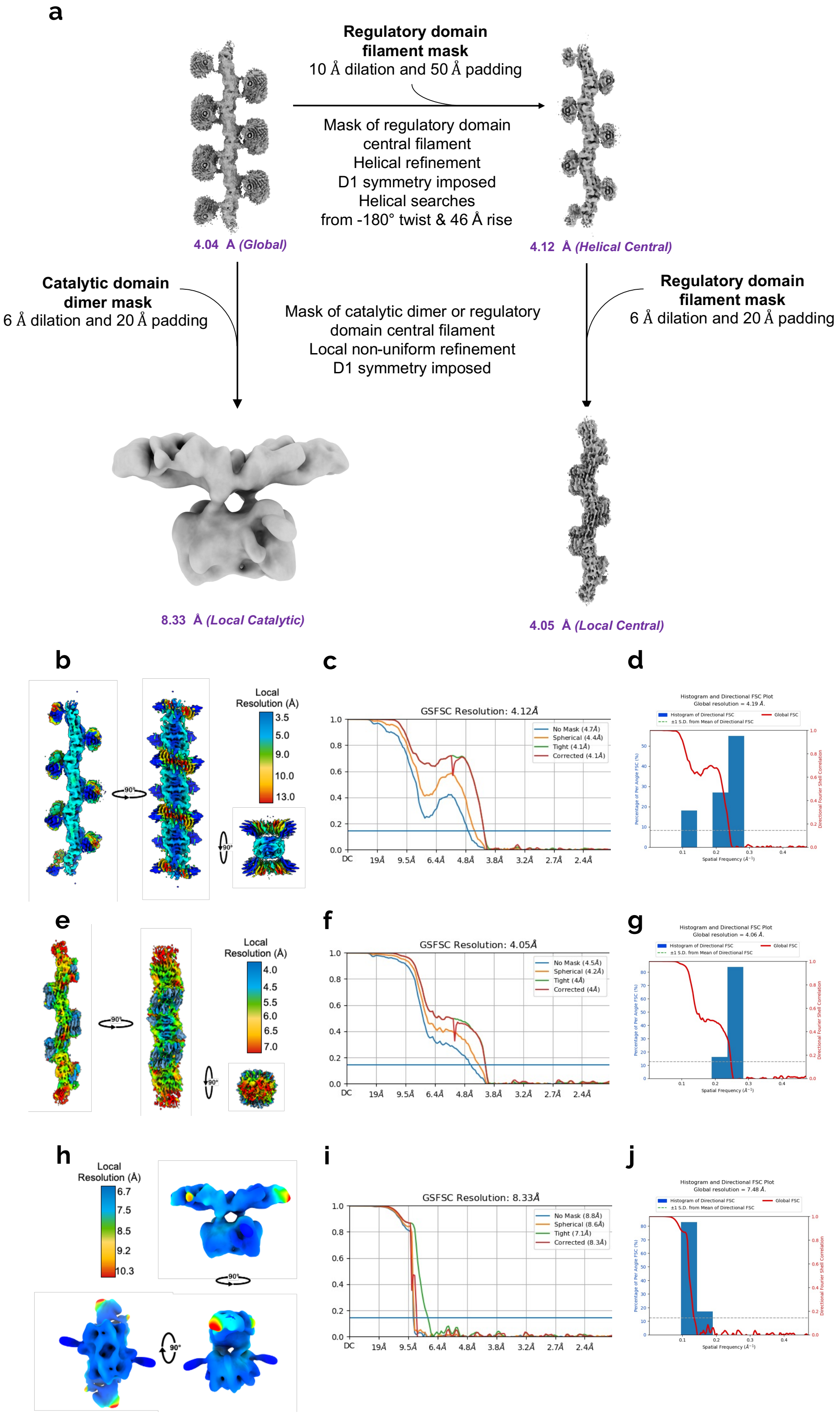


**Supplementary Fig. 6 | Further cryo-EM data processing of activated state CBS$^{FL\text{-}CHis}$. a**, Local helical and single particle processing flow chart of CBS$^{FL\text{-}CHis}$ in the activated, SAM bound, state. **b-d**, Local resolution, FSC curve, and directional FSC plot of the helical central map of CBS$^{FL\text{-}CHis}$ plus SAM. **e-g**, Local resolution, FSC curve, and directional FSC plot of the local central map of CBS$^{FL\text{-}CHis}$ plus SAM. **h-j**, Local resolution, FSC curve, and directional FSC plot of the local catalytic map of CBS$^{FL\text{-}CHis}$ plus SAM.

Source Images: Extended Data Figure 2

**a**

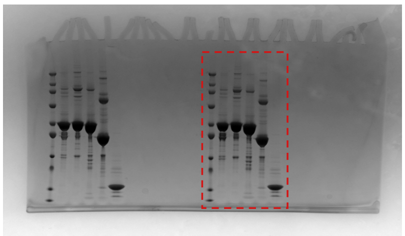

**b** $CBS^{FL}$

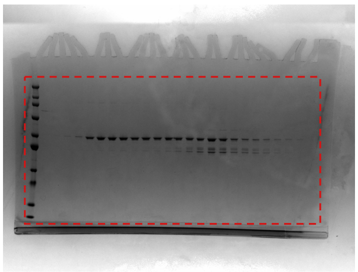

**b** $CBS^{FLC\text{-}His}$

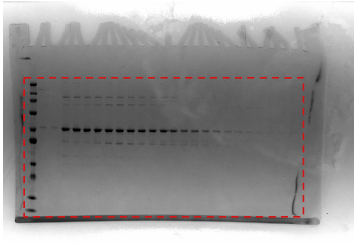

**b** $CBS^{\Delta 516\text{-}525}$

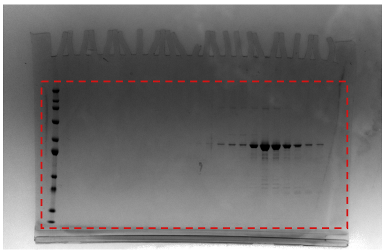

**b** $CBS^{CD}$

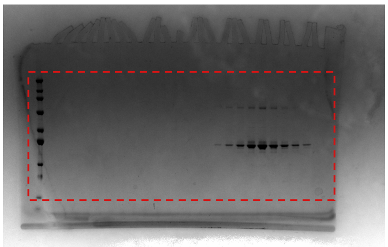

**b** $CBS^{RD\Delta 516\text{-}525}$

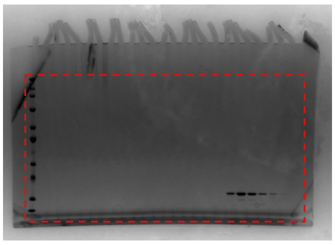

**c**

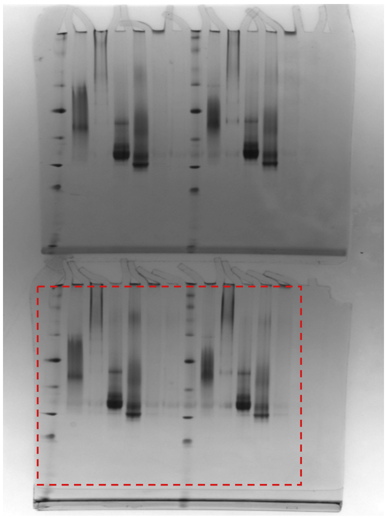

**d**

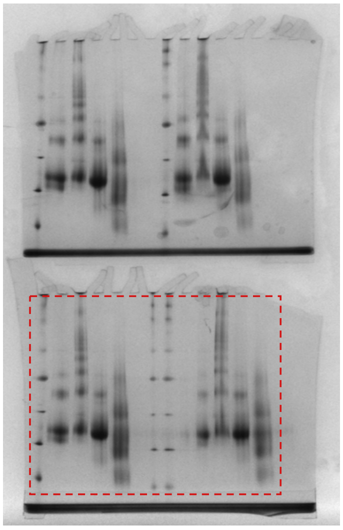

**Supplementary Fig. 7 | Source images of Extended Data Fig. 2a-d**, The dashed red box represents the cropped area used for the figure.

**Supplementary Movie 1 | Morph of the Bateman-Bateman interface from the basal and SAM bound activated structure of CBS.** The two neighbouring Bateman domains are represented as cartoons. SAM is represented as balls and sticks and is coloured pink.

**Supplementary Movie 2 | Morph of one full turn of the basal and SAM bound activated structures of CBS.** One CBS dimer is coloured light and dark blue. Neighbouring CBS dimers are coloured grey
